# Exploiting Endogenous CRISPR-Cas9 System for Functional Engineering of Probiotic *Lacticaseibacillus rhamnosus* GG (LGG)

**DOI:** 10.1101/2025.01.21.634116

**Authors:** Zifan Xie, Yong-Su Jin, Michael J. Miller

**Affiliations:** Department of Food Science and Human Nutrition, University of Illinois Urbana-Champaign, Urbana, IL, USA; Carl R. Woese Institute for Genomic Biology, University of Illinois Urbana-Champaign, Urbana, IL, USA

**Author notes:** Corresponding author: Prof. Yong-Su Jin Prof. Michael J. Miller.

**Keywords:** CRISPR-Cas9, *probiotic*, *synthetic biology*, *lactic acid bacteria*

## Abstract

*Lacticaseibacillus rhamnosus* GG (LGG) is one of the most extensively studied probiotic strains, widely used in food and health applications. However, the absence of efficient, precise genome editing methods has limited its broader potential and functional versatility. Here, we present an endogenous type II-A CRISPR-Cas9 genome editing workflow for LGG designed for functional strain construction. Using a plasmid interference assay together with single-nucleotide substitutions, we confirm the precise PAM requirement as 5′-NGAAA-3′. We pair a synthetic sgRNA cassette with homology-directed repair donors to enable precise deletions and insertions across multiple loci. Using this optimized and precise genome engineering method, we generated a β-glucuronidase (GUS)-expressing LGG strain for robust strain tracking within complex gut microbial communities. This work removes barriers to LGG engineering, expands the probiotic CRISPR toolkit, and provides broadly applicable strategies for designing next-generation probiotics with applications in food biotechnology and microbial therapeutics.

**GRAPHICAL ABSTRACT:** 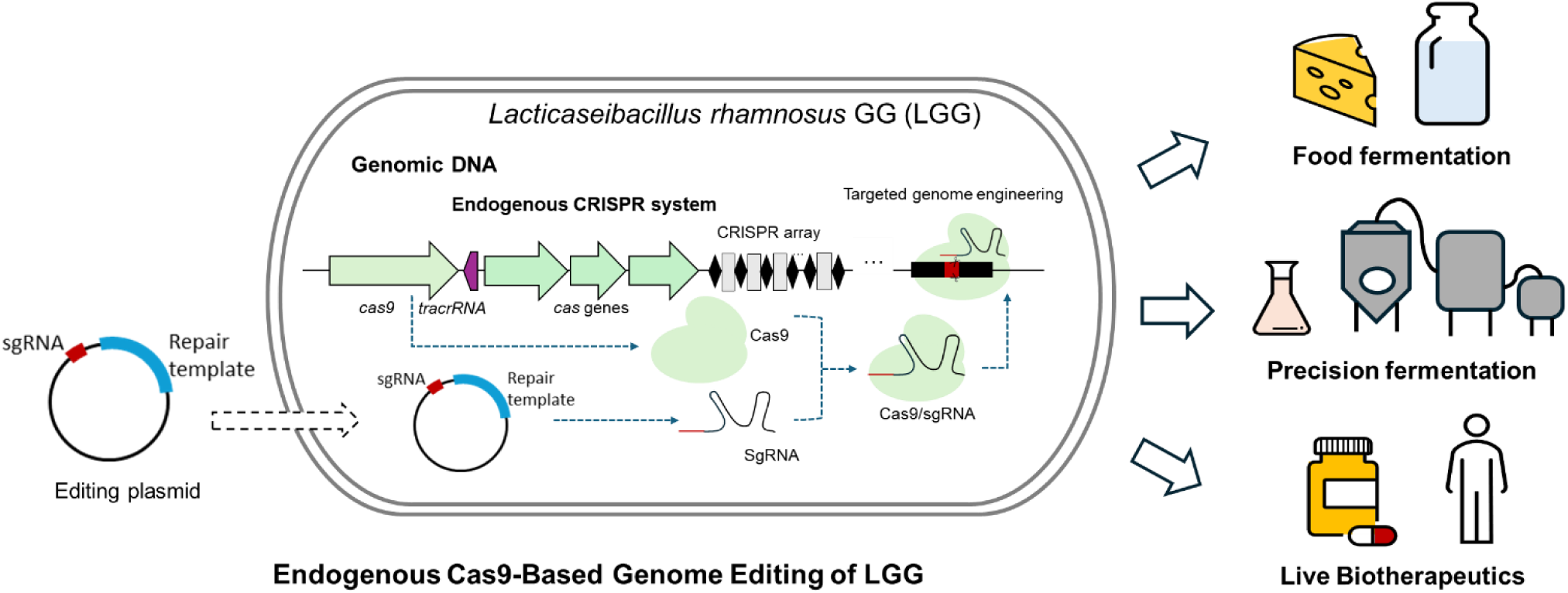

## INTRODUCTION

*Lacticaseibacillus rhamnosus* GG (LGG) is one of the most extensively studied probiotic strains due to its well-documented therapeutic and industrial benefits. Originally isolated from the human gastrointestinal tract, LGG demonstrates robust tolerance to gastric acid and bile salts, strong adherence to intestinal epithelial cells, and exceptional stability throughout food processing (Segers & Lebeer, 2014; Capurso, 2019). Since its discovery, numerous clinical and preclinical studies have demonstrated LGG’s ability to enhance gut barrier function, modulate host immune system, and exert antimicrobial activity (Hudault et al., 1997; Yan & Polk, 2002; Sindhu et al., 2014; Gao et al., 2019). Given its Generally Recognized as Safe (GRAS) status from the US Food and Drug Administration (GRAS Notice No. 845), LGG has been widely used in dietary supplements, fermented food products, and clinical therapies aimed at modulating gut health, immune modulation, and microbial homeostasis (Jia et al., 2016; Chan et al., 2021). More recently, LGG’s well-established safety profile and versatile functionality have attracted significant interest in its development as a microbial chassis for next-generation live biotherapeutics and precision fermentation applications (Günaydın et al., 2014; Wan et al., 2016; Petrova et al., 2018; Li et al., 2025).

Despite over 300 human-focused publications underscoring LGG’s extensive health benefits, its genetic manipulation and strain optimization have been significantly limited by a lack of efficient genome editing tools. Because of limited replicable plasmids, chromosomal modification in LGG has typically relied on non-replicative suicide vectors (Lebeer et al., 2012; S. Zhang et al., 2018). These approaches depend on low-frequency double-crossover events and usually require multiple rounds of selection or counterselection, making them labor-intensive, time-consuming, and poorly suited to complex or high-throughput engineering. The absence of precise genome-editing technologies has thus hindered mechanistic studies of LGG’s probiotic actions and restricted the development of advanced strains for therapeutic and industrial applications.

Recently, many innovative genome modification approaches have been developed and applied in lactic acid bacteria to determine relevant phenotypes and to enhance beneficial properties (Xie et al., 2024). Notably, the CRISPR-Cas (clustered regularly interspaced short palindromic repeats and CRISPR-associated proteins) system, originally discovered as a bacterial adaptive immune mechanism, has emerged as a powerful tool for precise and efficient genome modification in various organisms (Hsu et al., 2014; Adli, 2018). Heterologous CRISPR-Cas systems have been successfully employed in many lactobacilli, such as *Limosilactobacillus reuteri*, *Lactiplantibacillus plantarum*, *Lacticaseibacillus casei*, *Lactobacillus acidophilus*, *Lactobacillus gasseri*, and *Lacticaseibacillus paracasei* (Song et al., 2017; Leenay et al., 2018; Huang et al., 2019; Goh & Barrangou, 2021). Despite these successes, heterologous CRISPR systems present several challenges. These systems generally require at least one plasmid for the expression of large Cas9 protein, most commonly derived from *Streptococcus pyogenes* (Cong et al., 2013). Such large plasmid constructs often exhibit reduced plasmid stability, lower transformation efficiency, and impose metabolic burdens on host cells. Additionally, heterologous Cas proteins may exhibit suboptimal activity or cytotoxicity in certain strains, further limiting editing efficiency and viability (J. Zhang et al., 2018; Vento et al., 2019). To overcome these limitations, recent research has increasingly focused on endogenous CRISPR-Cas systems encoded within the host genomes. These native systems eliminate the need for exogenous proteins and exhibit enhanced compatibility with host machinery, resulting in improved editing precision and reduced cellular toxicity. Functional CRISPR-Cas systems have been identified in many lactobacilli strains (Crawley et al., 2018), and some of them have been successfully repurposed for precise genome editing. For example, the endogenous subtype I-E CRIPSR-Cas system has been utilized for efficient genetic manipulation of *Lactobacillus crispatus* (Hidalgo-Cantabrana, Goh, Pan, et al., 2019), and the subtype II-A CRISPR-Cas9 system has been used to achieve genome engineering in *Lcb. paracasei* (Gu et al., 2022). Recently, type II-A CRISPR-Cas9 system was identified in LGG (Crawley et al., 2018; Panahi et al., 2023), and the Cas9 protein from LGG has been heterologously expressed for plant genome engineering (Zhong et al., 2023). However, those studies did not establish a practical genome-editing workflow directly inside LGG.

Here, we establish an endogenous type II-A CRISPR-Cas9 genome-editing system for LGG that enables precise gene deletions and insertions. Using a plasmid-interference assay with single-nucleotide protospacer adjacent motif (PAM) variants, we confirmed the specificity of the native Cas9 PAM as 5′-NGAAA-3′. A replicative editing plasmid was constructed to carry synthetic single-guide RNA (sgRNA) cassettes and homologous repair templates, enabling efficient genome modifications across multiple loci. To demonstrate the practical applicability of this system, we engineered a β-glucuronidase (gusA) expressing strain that allows precise tracking in complex microbial communities and host models. Animal experiments further confirmed that the gusA-positive LGG strain serves as a suitable model for studying LGG-associated health benefits. Together, these results establish an efficient LGG genome-editing platform that facilitates mechanistic investigations and accelerates the development of functional probiotic strains for food biotechnology and microbial therapeutics.

## RESULTS

### Specificity of the Endogenous CRISPR-Cas9 System in LGG

LGG harbors a complete endogenous type II-A CRISPR-Cas9 locus containing four Cas protein-encoding genes (*cas1*, *cas2*, *csn2*, and *cas9*) adjacent to a CRISPR array (Crawley et al., 2018; Panahi et al., 2023). The array contains 24 distinct 30-base pair (bp) spacers interspaced by 36-bp direct repeats (Figure 1A). Consistent with previous reports, alignment of protospacer-flanking sequences in LGG indicates a preferred PAM of 5′-NGAAA-3′ (Figure 1B) (Crawley et al., 2018; Zhong et al., 2023)

**Figure 1.**
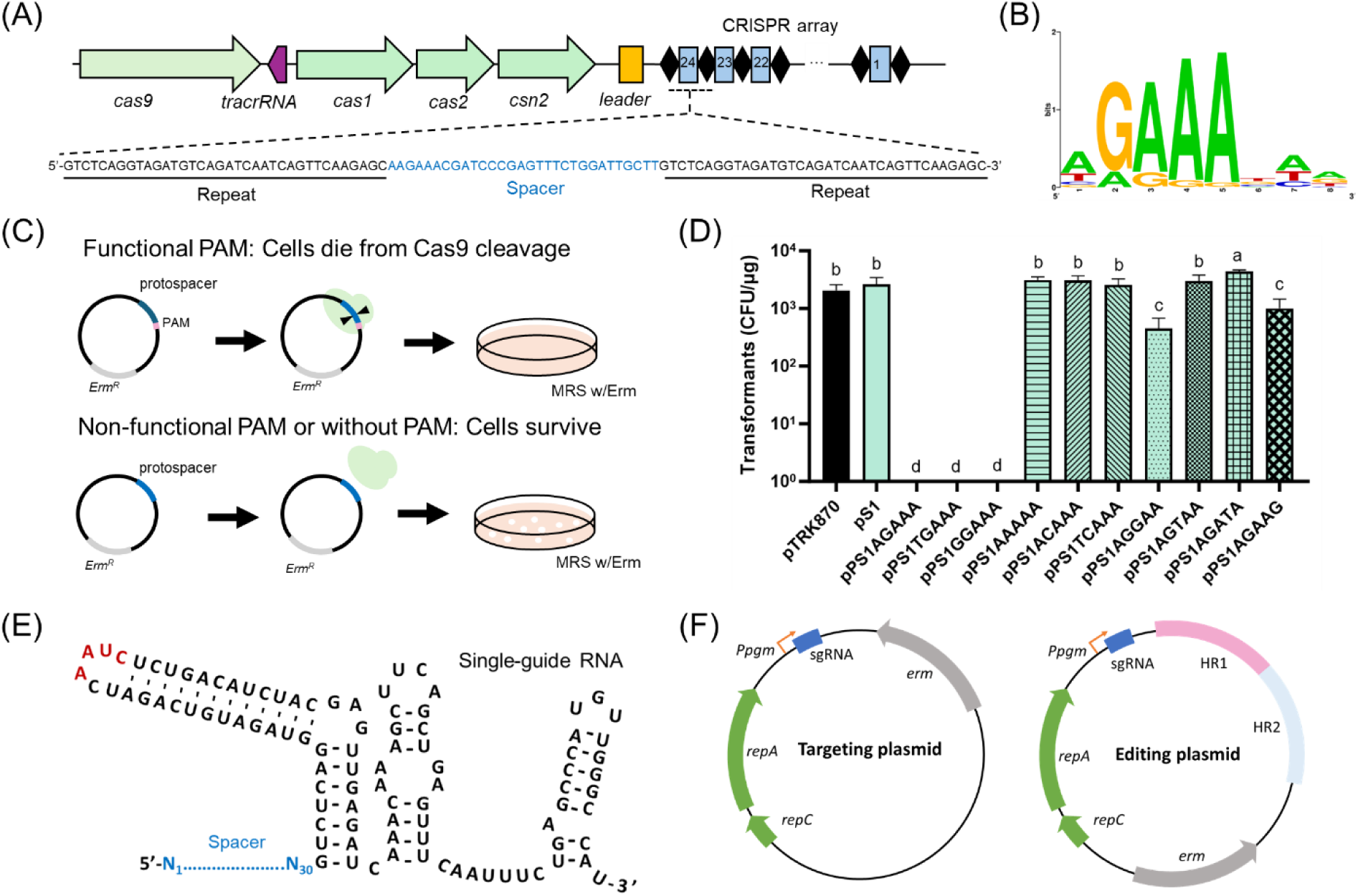
Genome-editing platform based on the endogenous type II-A CRISPR–Cas9 system in LGG. (A) Architecture of the CRISPR-Cas locus in LGG. The type II-A CRISPR-Cas9 system contains four Cas protein-encoding genes (*cas1, cas2, csn2*, and the critical *cas9*). The CRISPR array includes 24 spacers (30 bp, blue rectangles) interspaced with repeats (black diamonds). (B) PAM prediction visualized using a frequency plot generated using WebLogo. (C) Schematic of the plasmid interference assay. Plasmids carrying target sequences with functional or mutated PAMs were introduced into LGG to assess CRISPR-Cas9-mediated interference. A reduction in colony recovery indicates effective targeting by the native CRISPR-Cas9 system. (D) Recovery of transformants following introduction of interference plasmids compared to the control plasmid pTRK870. CFU: colony forming unit. Distinct letters were employed to denote significant differences (*p < 0.05*). (E) Structure of the SgRNA, designed by linking the crRNA and tracrRNA with an artificial AAUC loop (red). (F) Schematic of targeting and editing plasmids. *repA* and *repC*, plasmid replication gene; *erm*, erythromycin resistance gene; HR1 and HR2, the homologous repair templates; Ppgm, the promoter of phosphoglycerate mutase (*Ppgm*) from *L. acidophilus* NCFM.

To validate the specificity of the PAM sequence, we performed a plasmid interference assay incorporating single-nucleotide PAM substitutions (Figure 1C). Interference plasmids were constructed based on vector pTRK870 (Supplementary Table S2), a derivative of pTRK830 with a multiple cloning site (Xie et al., 2025). Each plasmid carried a protospacer matching the most recently acquired spacer 24 (5’-AAGAAACGATCCCGAGTTTCTGGATTGCTT-3’) in the CRISPR array, either accompanied by or lacking the predicted PAM sequence.

Transformation of LGG with plasmids (pPS1AGAAA, pPS1TGAAA, and pPS1GGAAA) containing the predicted PAM sequences resulted in no detectable transformants, demonstrating robust interference mediated by the endogenous CRISPR-Cas9 system (Figure 1D). To further evaluate PAM specificity, we introduced single-nucleotide substitutions at conserved positions (positions 2-5) of the predicted PAM sequence, generating seven PAM variants AAAAA, ACAAA, TCAAA, AGGAA, AGTAA, AGATA, and AGAAG. As shown in Figure 1D, plasmids with altered PAM sequences, pPS1AAAAA, pPS1ACAAA, and pPS1TCAAA, failed to produce a detectable interference response, indicating that guanine at position 2 is essential for effective PAM recognition. Additionally, plasmid pPS1AGATA, carrying a thymine substitution at position 4, showed significantly reduced interference, indicating the importance of adenine conservation at this position. Notably, plasmids pPS1AGGAA and pPS1AGAAG, each harboring substitutions at positions 3 and 5, exhibited partial interference, suggesting a moderate tolerance of these positions to guanine substitutions. However, significantly fewer colonies were recovered using the optimal PAM (pPS1AGAAA), confirming a strong preference for adenines at positions 3–5.

Collectively, these results confirm the PAM recognized by the endogenous Cas9 in LGG as 5′-NGAAA-3′ and demonstrate its high sequence specificity, with enhanced interference activity observed under our assay conditions.

### Construction of Editing Plasmids for Reprogramming the Endogenous CRIPSR-Cas9 System

After confirming the specificity of PAM sequence, we then explored its potential for genome editing in LGG. Initially, we designed an editing plasmid derived from pTRK870, which contains a homologous repair template and a CRISPR RNA (crRNA) expression cassette under control of the native CRISPR array leader sequence (Supplementary Figure S1). However, we found that due to the inherent instability of rolling circle plasmids (Xie et al., 2025), the space region between the repeats was frequently looped out from the plasmid during plasmid replication, resulting in spacer-deficient plasmids and a high frequency of wild-type (WT) survivors (Supplementary Figure S1).

To overcome this limitation, we transitioned to a sgRNA-based approach for directing precise genome targeting by the endogenous CRISPR-Cas9. Based on the predicted structure of the crRNA/ trans-activating CRISPR RNA (tracrRNA) duplex and prior transcriptional sequencing data (Crawley et al., 2018), we engineered an artificial sgRNA by linking the crRNA and tracrRNA with an AAUC loop (Figure 1E). We then constructed a targeting plasmid encoding the sgRNA under the control of the strong constitutive promoter *Ppgm* from *L. acidophilus* NCFM (Figure 1F) (Xie et al., 2025).

To validate our artificial sgRNA design, we targeted a highly conserved region within the 16S ribosomal RNA (rRNA) gene. Introduction of this 16S-targeting plasmid into LGG significantly reduced viable transformants compared to the pTRK870 control plasmid (Supplementary Figure S2), demonstrating effective Cas9-mediated chromosomal cleavage and providing direct evidence of functional sgRNA expression. To evaluate the broader applicability of this sgRNA system, we tested the same 16S-targeting plasmid in several additional *L. rhamnosus* strains (Supplementary Table S1). Unlike LGG, these other strains yielded viable transformants without evidence of Cas9-mediated lethality (Supplementary Figure S2), suggesting limited portability of our platform across closely related strains. Possible reasons include differences in plasmid transformation efficiency, strain-specific PAM sequence preferences, or variations in CRISPR architecture among strains (Supplementary Figure S2).

Building upon this validated targeting plasmid, we next designed a comprehensive editing plasmid incorporating both the sgRNA expression cassette and homologous repair templates (HR1 and HR2) flanking the intended genomic modification site (Figure 1F).

### Reprogramming the Endogenous CRISPR-Cas9 System for Gene Deletion

As a test, we first attempted to target the *fucI* gene (LGG_02685), which encodes L-fucose isomerase (Figure 2A). Deletion of *fucI* is expected to significantly impair LGG growth when fucose is the sole carbon source (Becerra et al., 2015). A 30-bp DNA sequence complementary to *fucI* was selected as the spacer and cloned into the pTRK870 backbone, together with two 1-kb homologous arms, to generate the editing plasmid pfucI-sgRNA-RT (Supplementary Table S2). As controls, we also constructed plasmids containing only the repair template (pfucI-RT) or only the sgRNA (pfucI-sgRNA) (Supplementary Table S2). Transformation of LGG with the self-targeting plasmid (pfucI-sgRNA) yielded no colonies (Figure 2B), confirming effective genome cleavage triggered by the endogenous CRISPR-Cas9 system guided by the sgRNA. In contrast, transformation with the repair template-only plasmid (pfucI-RT) generated numerous colonies (Figure 2B), but none of screened colonies carried the intended deletion genotype (Supplementary Figure S3), indicating that the repair template alone was insufficient for generating targeted deletions efficiently. Importantly, transformation with the complete editing plasmid (pfucI-sgRNA-RT) resulted in significantly fewer colonies (four colonies). PCR screening of these transformants identified one colony (Colony No. 1) displaying the expected 2.1-kb PCR product, indicative of the deletion genotype, whereas the other three exhibited wild-type PCR products (2.7 kb; Figure 2C). Sanger sequencing of the 2.1-kb amplicon confirmed the precise deletion, and the successfully engineered strain was designated MJM568 (Figure 2D). Phenotypic analysis further revealed that the MJM568 strain could not utilize fucose as the sole sugar source (Figure 2E).

**Figure 2.**
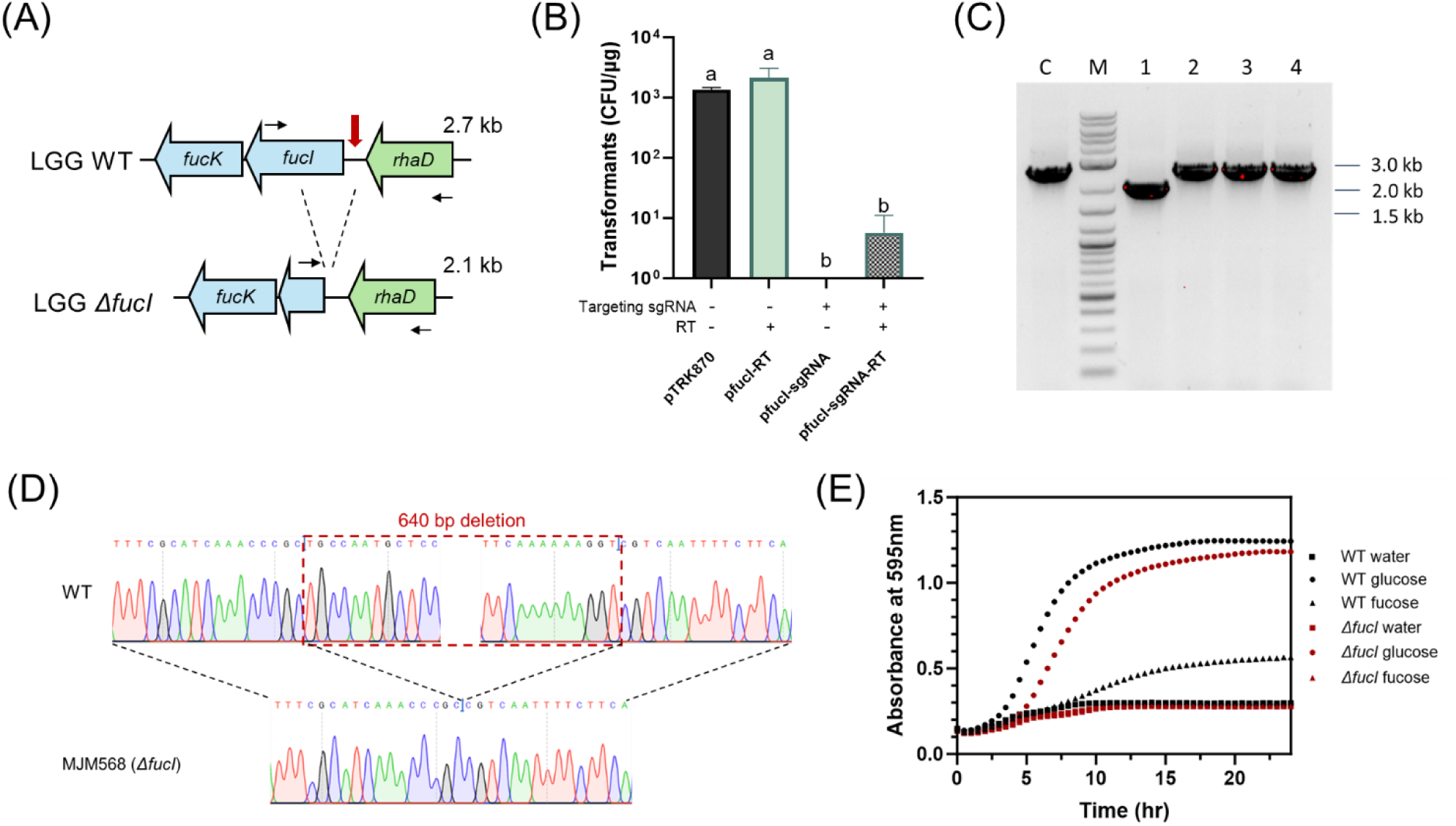
Deletion of L-fucose isomerase (*fucI*) by reprogramming the endogenous CRISPR-Cas9 system in LGG. (A) Schematic representation of the genotypes of the wild type (WT) strain and the *ΔfucI* mutant. (B) Transformation efficiencies of control plasmid pTRK870, homologous repair template containing plasmid pfucI-RT, self-targeting plasmid pfucI-sgRNA, and editing plasmid pfucI-sgRNA-RT. Distinct letters were employed to denote significant differences (*p < 0.05*). (C) Colony PCR screening of four surviving transformants carrying the editing plasmid pfucI-sgRNA-RT to confirm the deletion genotype. Expected PCR product sizes for WT and *ΔfucI* are 2.7 kb and 2.1 kb, respectively. C: WT control; M: DNA marker; 1-4: surviving transformants. (D) Sanger sequencing results confirming the 640-bp deletion in the Δ*fucI* strain (MJM568) compared to WT. (E) Growth curve of WT (black) and MJM568 (red) strains in MRS broth containing water (square), 2% glucose (circle) or 2% fucose (triangle), measured by absorbance at 595nm.

To further evaluate the versatility of this system, we successfully deleted a larger 1.5-kb genomic region encoding coenzyme A (CoA) synthase (*acs1*) using the same strategy (Supplementary Figure S4). This demonstrated the platform’s capability for targeting and deleting genomic regions of varying lengths. Collectively, these results confirm that the LGG endogenous CRISPR-Cas9 system can be effectively harnessed for targeted gene deletions, with both the sgRNA-guided Cas9 cleavage and homologous repair templates essential for achieving precise genomic modifications.

### Reprogramming the Endogenous CRISPR-Cas9 System for Gene Insertion

After demonstrating the feasibility of gene deletions using the native CRISPR-Cas9 system in LGG, we next aimed to evaluate the system’s capacity for gene insertion. The ability to stably integrate heterologous genes is a critical feature for engineering LGG as a microbial cell factory or live biotherapeutic. However, the previously used plasmid backbone pTRK870, which replicates via a rolling-circle mechanism, is inherently unstable and has limited insert capacity (Xie et al., 2025). To address these limitations, we constructed a new editing backbone, pMJM100 (Supplementary Figure S5), which is a smaller and more stable plasmid with improved capacity for accommodating larger genetic inserts. This redesigned backbone provides a robust platform for gene insertion mediated by the native CRISPR-Cas9 machinery in LGG.

To test this platform, we repurposed the *fucI*-targeting sgRNA expression cassette previously used for gene deletion and combined it with a homologous repair template designed to partially delete *fucI* and simultaneously insert a chloramphenicol resistance gene (*cam*) expression cassette (Figure 3A). The *cam* cassette was flanked by 0.5 kb homology arms derived from the original *fucI* deletion template. The use of shorter arms reduced the plasmid size and improved its stability during cloning and transformation. The resulting construct, pfucI-sgRNA-Cm-RT2, was assembled using the pMJM100 backbone.

**Figure 3.**
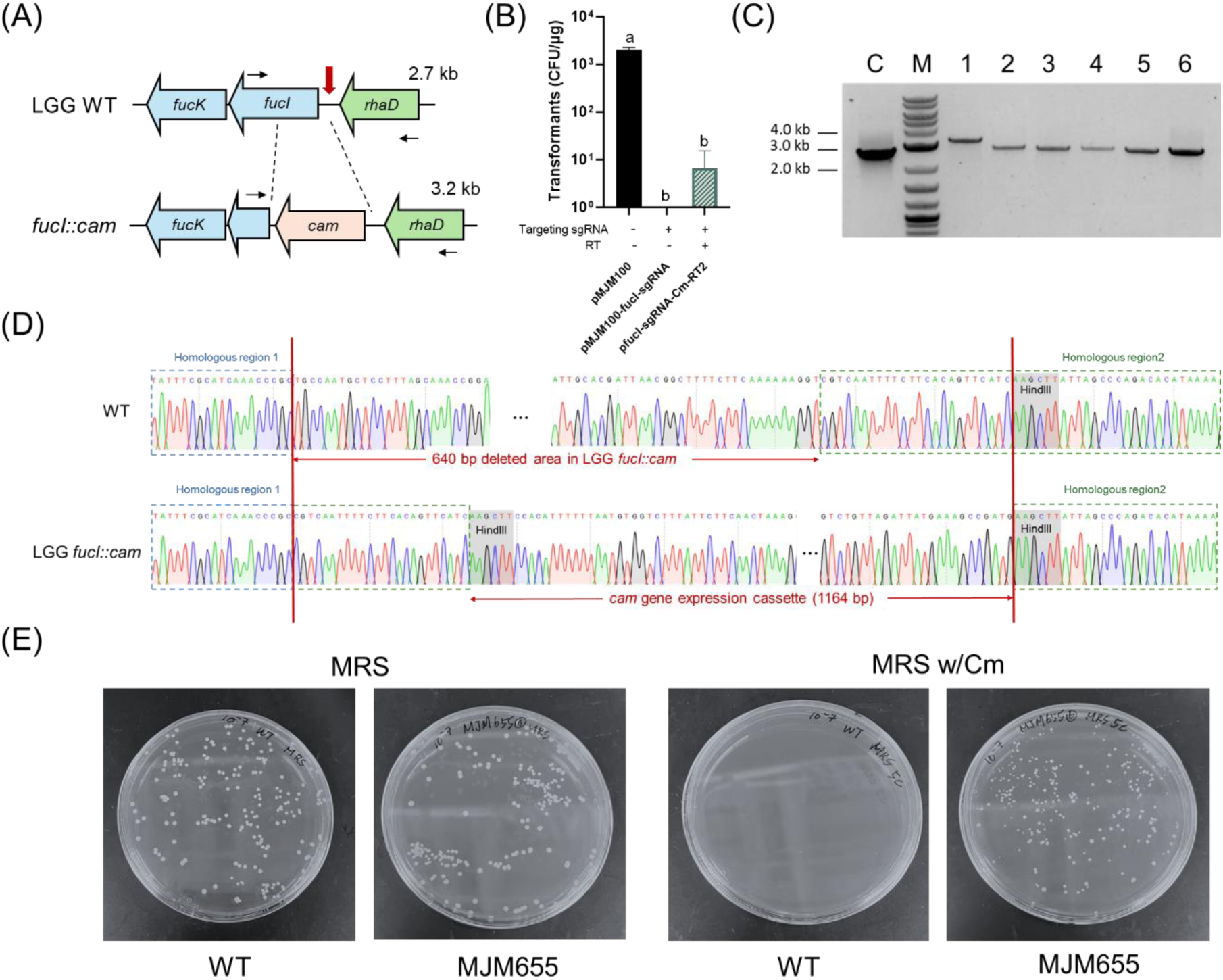
Insertion of the chloramphenicol resistance (*cam*) gene by reprogramming the endogenous CRISPR-Cas9 system in LGG. (A) Schematic representation of the genotypes of the wild type (WT) strain and the LGG *fucI::cam* mutant. (B) Transformation efficiencies of control plasmid pMJM100, self-targeting plasmid pMJM100-fucI-sgRNA, and editing plasmid pfucI-sgRNA-Cm-RT2. Distinct letters were employed to denote significant differences (*p < 0.05*). (C) Colony PCR screening of six surviving transformants carrying the editing plasmid pfucI-sgRNA-Cm-RT2 to confirm the insertion genotype. Expected PCR product sizes for WT and *fucI::cam* are 2.7 kb and 3.2 kb, respectively. C: WT control; M: DNA marker; 1-6: surviving transformants. (D) Sanger sequencing results of the WT and LGG *fucI::cam* strains. In *fucI::cam*, a 1164-bp *cam* gene expression cassette was inserted into the genome, replacing a 640-bp segment of the *fucI* gene. Homologous region 1 is enclosed in a dashed blue box, and homologous region 2 is enclosed in a dashed green box. The *HindIII* restriction site, located at the junction, is shaded in grey. (E) Growth of WT LGG and MJM655 strains on MRS plates or MRS plates supplemented with 5 µg/mL chloramphenicol.

Following transformation of pfucI-sgRNA-Cm-RT2 into LGG, six erythromycin-resistant transformants were obtained (Figure 3B). PCR screening revealed that one transformant (Colony No. 1) displayed the expected amplicon size for the *cam* gene insertion (Figure 3C), which was further confirmed by Sanger sequencing (Figure 3D). To eliminate the editing plasmid, the verified mutant was passaged twice under non-selective conditions, yielding an erythromycin-sensitive derivative (Supplementary Figure S6), designated MJM655. As excepted, MJM655 showed robust growth on MRS plates supplemented with 5 µg/mL chloramphenicol, while the WT LGG strain remained chloramphenicol-sensitive (Figure 3E).

### Construction of GUS-Positive LGG

To demonstrate the application of our endogenous CRISPR-Cas9 system for functional gene insertion, we engineered a GUS-positive LGG strain by inserting the *gusA* gene from *Lactobacillus gasseri* ADH, a GRAS strain. The *gusA* gene encodes β-glucuronidase and is widely used as a reporter in lactic acid bacteria. We aimed to create a chromosomally integrated reporter strain suitable for tracking in complex microbial communities.

To achieve this, we constructed the editing plasmid prha-sgRNA-gusA-RT2, designed to introduce *gusA* gene expression cassette into the intergenic region between the fucose and rhamnose operons (Figure 4A). Following transformation and selection, colony PCR of one transformant (Colony No. 1) showed both wild-type and mutant bands (Figure 4B), indicating a mixed population. Re-isolation of colony No.1 yielded colonies with a single PCR product of the expected insertion size (Supplementary Figure S7), and Sanger sequencing confirmed precise genomic integration of the *gusA* cassette at the target locus (Figure 4C). After plasmid curing through non-selective subculturing, the resulting genome-edited strain was designated MJM670. To verify *gusA* expression, we plated MJM670 and WT LGG on MRS agar supplemented with X-gluc. As expected, MJM670 formed distinct blue colonies, while the WT LGG colonies remained white (Figure 4D), confirming successful chromosomal expression of *gusA*. Moreover, MJM670 maintained consistent GUS expression over serial passages, in contrast to a plasmid-based *gusA* construct, which showed progressive loss of expression (Figure 4E), underscoring the advantage of chromosomal integration for long-term genetic stability.

**Figure 4.**
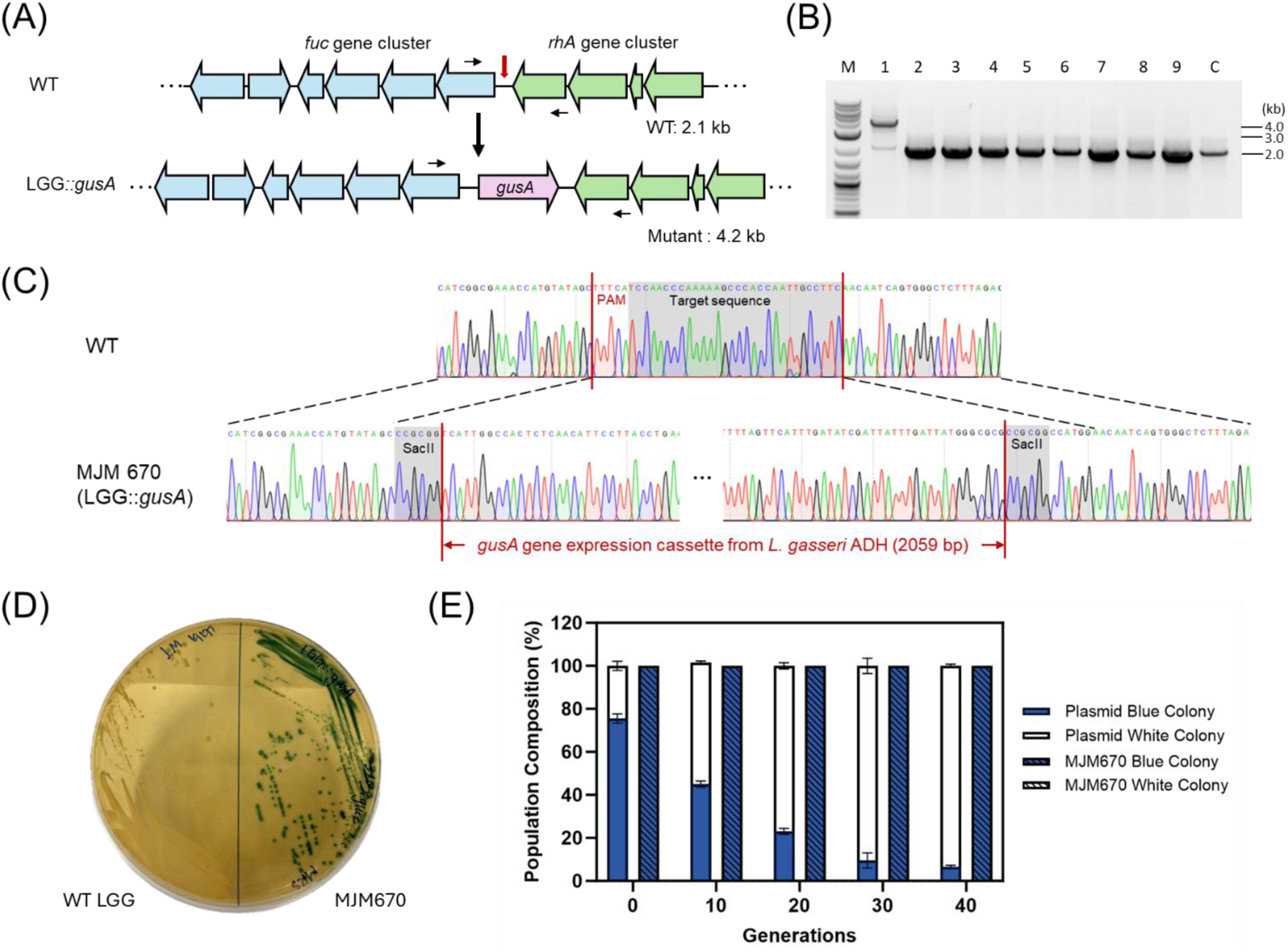
Insertion of the β-glucuronidase encoding gene *gusA* by reprogramming the endogenous CRISPR-Cas9 system in LGG. (A) Schematic representation of the genomic locus in the wild type (WT) strain and the LGG::*gusA* mutant, showing insertion of the *gusA* expression cassette from *Lactobaillus gasseri* ADH between the fucose and rhamnose operons. (B) Colony PCR screening of nine surviving transformants carrying the editing plasmid prha-sgRNA-gusA-RT2 to confirm the insertion genotype. Expected PCR product sizes for WT and LGG::*gusA* are 2.1 kb and 4.2 kb, respectively. C: WT control; M: DNA marker; 1-9: surviving transformants. (C) Sanger sequencing results confirming precise *gusA* insertion in the mutant strain MJM670 (LGG::*gusA*) compared to WT. (D) Growth phenotype of WT LGG and MJM670 on MRS agar supplemented with X-gluc. MJM670 exhibits a blue colony phenotype indicative of β-glucuronidase activity, while WT colonies remain white. (E) Stability of *gusA* gene expression across generations for plasmid-based and genome-integrated constructs.

To evaluate whether the insertion affected adjacent gene function, we compared the growth profiles of MJM670 and WT LGG in media supplemented with glucose, fucose or rhamnose. As shown in Supplementary Figure S8, MJM670 exhibited growth patterns nearly identical to the wild-type strain under all tested sugar conditions, indicating that *gusA* integration did not disrupt the function of the flanking fucose and rhamnose operons. Together, these results demonstrate that the intergenic region between the fucose and rhamnose operons is a permissive site for gene insertion, and that prha-sgRNA-RT2 is an effective vector for functional gene integration in LGG. This strategy offers a valuable tool for constructing recombinant probiotic strains for metabolic enhancement and therapeutic applications.

### Quantification of GUS-Positive LGG in Mouse Fecal Samples

Analyzing LGG abundance in fecal samples offers a non-invasive and comprehensive view of bacterial persistence through the gastrointestinal (GI) tract. Fecal enumeration is particularly valuable for assessing long-term colonization and microbial dynamics, providing insight into the overall stability and retention of probiotic strains in vivo. To evaluate the persistence and viability of engineered GUS-positive LGG, MJM670, we conducted both plating-and qPCR-based analysis of fecal samples collected from C57BL/6J mice following a single oral gavage (Figure 5A).

**Figure 5.**
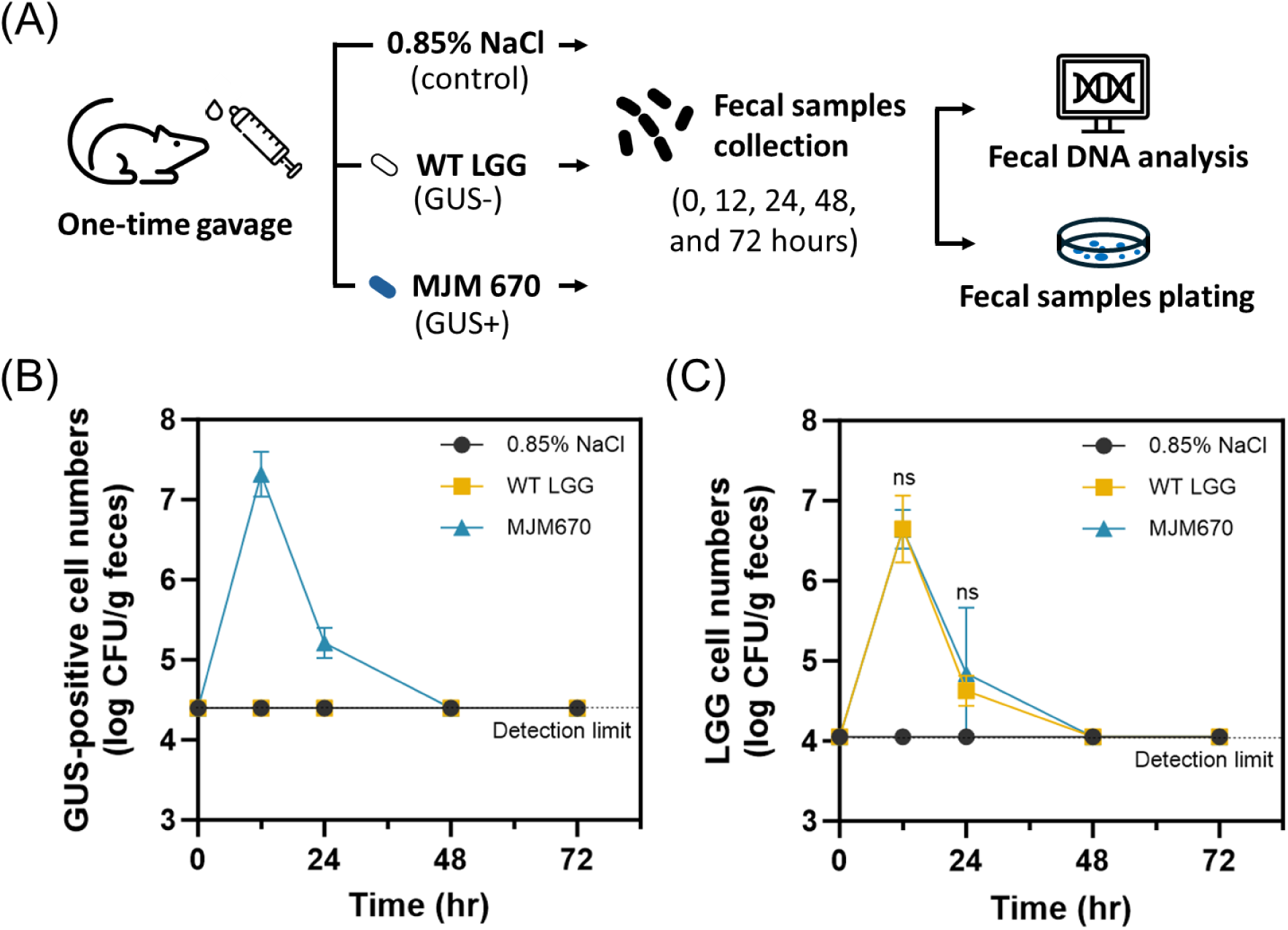
*In vivo* tracking and quantification of GUS-positive LGG in mice. (A) Mice were orally gavaged with saline, WT LGG (GUS-LGG), or MJM670 (GUS+ LGG). Fecal samples were collected at 0, 12-, 24-, 48-, and 72-hours post-gavage. (B) MJM670 was quantified by plating on MRS + X-gluc agar to enumerate blue colonies (viable GUS+ LGG). (C) In parallel, total LGG abundance was assessed in all groups by LGG-specific qPCR. Data represents mean ± SD from biological replicates (n=6).

As previously shown in Figure 4D, MJM670 expresses β-glucuronidase, enabling colony visualization on MRS agar supplemented with X-Gluc. GUS activity resulted in distinct blue colonies, allowing clear differentiation from native gut microbiota (Supplementary Figure S9). Based on plating analysis (Figure 5B), approximately 10⁷ CFU/g MJM670 were detected in feces 12 hours after one-time gavage, which decreased to ∼10⁵ CFU/g at 24 hours. By 48 hours, MJM670 was undetectable in all groups, indicating complete clearance. In parallel, we performed qPCR using LGG-specific primers to assess total LGG abundance regardless of cell viability (Figure 5C). The qPCR results revealed that MJM670 exhibited similar survival dynamics in the GI tract as the WT LGG strain (Figure 5C), with both showing a rapid decline over time. These results further support the observation that both WT LGG and MJM670 exhibit transient colonization following oral administration.

When comparing the two methods (Figure 5B and 5C), the plating method consistently yielded higher LGG counts than qPCR. This is likely due to incomplete DNA recovery from fecal samples using commercial stool DNA kits, which may underestimate total bacterial abundance. Moreover, while qPCR can detect both viable and nonviable cells, plating selectively recovers only viable LGG cells capable of colony formation, providing a direct measure of functional probiotic survival. Despite its higher detection threshold, the plating method provides unique advantages for engineered strains like MJM670. Its visual specificity enables clear identification of the target strain, and it allows isolation of individual colonies for downstream phenotypic or genomic characterization, which is an essential step for verifying genetic stability or strain behavior after going through the GI tract.

Overall, the GUS-positive LGG strain, MJM670, demonstrated reliable detection via plating and enabled viable cell recovery, supporting its application in colonization tracking and functional studies within the gastrointestinal microbiome.

## DISCUSSION

LGG is one of the most widely used probiotics in fermented foods and dietary supplements, yet the development of enhanced LGG strains has been constrained by the lack of efficient genome editing tools. Existing methods based on non-replicable vectors are inefficient, labor-intensive, and require antibiotic selection. While heterologous CRISPR-Cas9 systems have been applied in other *Lactobacillus* species, their use in LGG has been restricted due to the lack of stable plasmids capable of accommodating the large *cas9* gene. In this situation, repurposing the endogenous CRISPR–Cas machinery offers a promising approach for editing in LGG. A type II-A CRISPR–Cas9 system has previously been identified in LGG, and its Cas9 protein has been heterologously expressed for plant genome engineering (Crawley et al., 2018; Zhong et al., 2023). However, these studies did not establish a practical genome-editing workflow in LGG. Critical parameters, such as PAM sequence specificity in the LGG context, sgRNA expression system, homologous repair templates design, plasmid curing, and edit stability, have remained undefined. Here, we addressed these gaps and developed an endogenous type II-A CRISPR-Cas9 platform for LGG that enables efficient genome editing to construct functional strains for fermentation and therapeutic applications.

We first confirmed 5’-NGAAA-3’ as the preferred PAM, exhibiting strict sequence requirements for target recognition. To assess the reprogrammability of the native system, we designed an artificial sgRNA targeting the essential 16S rRNA gene. Introduction of this construct resulted in a marked loss of viable transformants in LGG, confirming the high activity and specificity of the native system and demonstrating that artificial sgRNAs can be correctly recognized and processed in LGG. In contrast, the same construct failed to induce lethality in other *Lcb. rhamnosus* strains, despite some encoding type II-A CRISPR loci. These strains showed no detectable targeting activity, likely due to low transformation efficiency or variations in PAM specificity or CRISPR locus architecture. These findings highlight the strain-dependent nature of endogenous CRISPR systems and underscore the importance of strain-specific validation for successful repurposing.

Using the validated system, we developed replicable plasmids for sgRNA expression and repair template delivery, achieving precise deletions and insertions without scars or antibiotic-resistant genes, making them suitable for food and therapeutic applications. Although overall editing efficiency in LGG was lower than in some other lactobacilli (Gu et al., 2022; Hidalgo-Cantabrana, Goh, Pan, et al., 2019), likely due to the inherent instability of rolling-circle plasmids and relatively lower homologous recombination efficiency, this workflow offers several practical advantages. The system enables scarless gene deletions and insertions using a single plasmid, without the need for heterologous Cas9 expression or laborious selection. In practice, desired mutants can be routinely obtained in one transformation and a brief colony screen, typically completed within three days, without multiple rounds of subculturing.

Moreover, the editing plasmids are easily curable, leaving no selection or counterselection markers in the final strains, which support iterative marker-free editing to stack multiple modifications in the same genetic background. Together, these attributes make the platform well-suited for functional genetics, strain optimization, and medium-throughput genome engineering in LGG, where speed, precision, and regulatory compatibility are at a premium.

To demonstrate the application of this genome editing platform, we engineered a GUS-expressing LGG strain (MJM670) to enable specific and robust tracking within mixed microbial populations and animal models. In the gastrointestinal tract, endogenous β-glucuronidase activity is present in only a limited number of bacterial species beyond *Escherichia coli*, making *gusA* a reliable and specific reporter gene. Genomic integration into LGG enables clear visual differentiation from native gut microbes and provides a direct alternative to qPCR-based detection. While qPCR detects DNA from both viable and nonviable cells, plating recovers only viable LGG, providing a direct measure of functional probiotic survival. The *gusA* gene, sourced from the GRAS strain *L. gasseri* ADH, supports biosafety and avoids antibiotic resistance genes, reducing the risk of horizontal gene transfer and the spread of antibiotic resistance within the gut environment. This GUS-positive LGG enables both qualitative and quantitative monitoring in complex communities, like co-culture systems, fecal samples, and host tissues. Therefore, these features make GUS-positive LGG a valuable tool for mechanistic probiotic research and a promising chassis for therapeutic protein or antigen delivery in live biotherapeutics.

In conclusion, we present a functional genome editing platform for LGG based on its endogenous CRISPR-Cas9 system. This system enables precise, scarless gene deletions and insertions using a single plasmid. The platform is rapid, convenient, and broadly applicable, making it a valuable tool for LGG functional studies, probiotic engineering, and industrial strain development. The successful engineering of GUS-expressing LGG strains highlights the potential of this system for live biotherapeutic applications. In the future, this platform has the potential to broaden LGG’s role as a programmable chassis for therapeutic delivery and precision fermentation. By enabling efficient and scarless genome editing, our platform paves the way for future research and strain development in LGG and related *Lactobacillus* species, offering a powerful tool to enhance probiotic functionality and industrial utility.

## MATERIAL AND METHODS

### Strains and Culture Conditions

All *Lcb. rhamnosus* strains (listed in Supplementary Table S1) were cultivated in de-Man-Rogosa-Sharpe (MRS; Hardy Diagnostics, Santa Maria, CA, USA) broth or on MRS agar (1.5%) at 37 ℃ under anaerobic conditions (90% N_2_, 5% CO_2_, and 5% H_2_). *Escherichia coli* MC1061 was cultured in Luria-Bertani (LB) broth with agitation at 250 rpm or on LB agar (1.5%) and incubated at 37 ℃. For GUS activity detection, MRS agar plates were supplemented with 200 µg/mL 5-Bromo-4-chloro-1*H*-indol-3-yl β-D-glucopyranosiduronic acid (X-gluc; Gold Biotechnology, USA). When appropriate, antibiotics were added at the following concentrations: erythromycin (Em; Gold Biotechnology, St. Louis, MO, USA) at 5 µg/ml for LGG and 150 µg/ml for *E. coli*, and chloramphenicol (Cm; Sigma, St. Louis, MO, USA) at 5 µg/ml for LGG.

### CRISPR-Cas System Characterization *in silico*

The complete genome sequence of LGG (GenBank: CP031290.1) is available at the National Center for Biotechnology Information (NCBI) database. CRISPR array and associated Cas proteins were identified using the CRISPRCasFinder platform (Couvin et al., 2018). PAM sequences were predicted by aligning LGG spacer to target sequences with CRISPRTarget (Biswas et al., 2013), and consensus PAM sequences were presented as sequence logo generated by WebLogo (Crooks et al., 2004). For type II CRISPR systems, tracrRNA contains a region complementary to the repeat sequence of the crRNA; thus, the repeat sequence was queried against the genome using the Basic Local Alignment Search Tool (BLAST) to locate the tracrRNA (Hidalgo-Cantabrana, Goh, & Barrangou, 2019). The crRNA– tracrRNA duplex was further modeled using the NCPACK web server and manually illustrated (Zadeh et al., 2011).

### DNA Manipulations and Molecular Cloning

Genomic DNA was isolated using the Wizard genomic DNA purification kit (Promega Corporation, Madison, WI, USA). Plasmid DNA from *E. coli* was prepared with the QIAGEN plasmid mini-prep kit (QIAGEN, Germantown, MD, USA). Restriction enzymes and T4 DNA ligase ( New England Biolabs, Ipswich, MA, USA) were used according to the supplier’s protocols. Polymerase chain reactions (PCR) were conducted following standard procedures with either DreamTaq DNA polymerase (Thermo Fisher Scientific, Waltham, MA, USA) or Phusion DNA polymerase (NEB, Ipswich, MA, USA). Oligonucleotide primers (Supplementary Table S3) were ordered from Integrated DNA Technologies (IDT, Coralville, IA, USA) and synthetic gene fragments (Supplementary Table S4) were obtained from Twist Bioscience (Twist Bioscience, San Francisco, CA, USA). PCR amplicons were electrophoresed on 1.0% agarose gels and purified using the QIAquick PCR purification kit (QIAGEN, Germantown, MD, USA). Sanger sequencing was performed at the UIUC Core Sequencing Facility (Urbana, IL, USA), and whole plasmid sequencing was conducted by Plasmidsaurus, an Oxford Nanopore certified service.

### Construction of Interference Plasmids

pTRK830 is a replicating shuttle vector for *E. coli* and lactobacilli strains (Xie et al., 2025). The multiple cloning site (MCS) from pBluescript KS (+) was ligated into the *StuI* site of pTRK830, which generated pTRK870 (Supplementary Table S2). Plasmid pTRK870 was used for all following plasmid constructions. The interference plasmids were constructed as described before (Crawley et al., 2018).

The double-stranded DNA protospacers, with or without the PAM, were generated by PCR using and then ligated into XbaI-HindIII-digested pTRK870 to construct interference plasmids. The resulting interference plasmids were PCR-screened in *E. coli* transformants using PS1 and PS2 primers (Supplementary Table S3) for the presence of the insert and sequenced to confirm sequence content.

### Construction of Editing Plasmids

Each editing plasmid consisted of two key components: a sgRNA expression cassette and a repair template. The sgRNA expression cassettes included a promoter *Ppgm* of the phosphoglycerate mutase from *L. acidophilus* NCFM, a designed sgRNA containing a 30-bp sequence targeting a specific genomic locus of LGG strain, and a Rho-independent terminator.

#### *fucI* gene deletion

The *fucI* targeting sgRNA expression cassette (Supplementary Table S4) was synthesized and amplified using primers sgRNA-XbaI and sgRNA-XmaI (Supplementary Table S3), then cloned into XbaI-XmaI-digested pTRK870 to generate targeting plasmid pfucI-sgRNA (Supplementary Table S2). The repair template with two 1-kb homologous regions for *fucI* gene editing was amplified from the genomic DNA of LGG strain. For this, two 1-kb homologous regions were fused by overlap extension PCR to obtain the 2-kb repair template to the *fucI* gene. This repair template was digested with SalI and ApaI, then cloned into SalI-ApaI-digested plasmids pfucI-sgRNA and pTRK870 to generate plasmids pfucI-sgRNA-RT and pfucI-RT (Supplementary Table S2), respectively. All these final plasmid constructs were PCR-screened using PS1 and PS2 primers (pTRK870 based) and sequenced to confirm sequence content.

#### *acs1* gene deletion

The *acs1* targeting sgRNA expression cassettes (Supplementary Table S4) were synthesized and amplified using primers sgRNA-XbaI and sgRNA-XmaI (Supplementary Table S3), then cloned into XbaI-XmaI-digested pTRK870 to generate targeting plasmids pacs1-sgRNA (Supplementary Table S2). The repair template (Supplementary Table S4) with two 1-kb homologous regions was synthesized and cloned into the PstI-ApaI-digested pacs1-sgRNA generating pacs1-sgRNA-RT (Supplementary Table S2). All these final plasmid constructs were PCR-screened using PS1 and PS2 primers (pTRK870 based) and sequenced to confirm sequence content.

#### *cam* expression cassette insertion

To construct a compact and stable cloning vector for gene insertion, a multiple cloning site sequence was inserted into StuI-MfeI-digested pTRK829 (Supplementary Table S2), removing the entire *cam* gene and other non-essential regions, to generate pMJM100 (Supplementary Figure S2). The *fucI* targeting sgRNA expression cassette (Supplementary Table S4) was cloned into XbaI-XmaI-digested pMJM100 to construct a new *fucI* targeting plasmid pMJM100-fucI-sgRNA (Supplementary Table S2). The repair template, containing two 0.5-kb homologous regions, was amplified from pfucI-RT using primers fucI-RT2-SalI and fucI-RT2-ApaI (Table S3), and cloned into a TOPO vector using the Zero Blunt® TOPO® PCR Cloning Kit (Thermo Fisher Scientific, Waltham, MA, USA), resulting in pTOPO-fucI-RT2 (Supplementary Table S2). The *cam* gene expression cassette was amplified from pTRK829 using primers Cm-F and Cm-R and inserted into pTOPO-fucI-RT2 at the *HindIII* site, generating pTOPO-fucI-Cm-RT2 (Supplementary Table S2). Finally, the complete repair template, containing the *cam* gene expression cassette flanked by two 0.5-kb homologous regions, was cloned into SalI-ApaI-digested pMJM100-fucI-sgRNA, resulting in the final editing plasmid pfucI-sgRNA-Cm-RT2 (Supplementary Table S2). pfucI-sgRNA-Cm-RT2 was PCR-screened using PS3 and PS4 primers (pMJM100 based) and sequenced to confirm sequence content.

#### *gusA* expression cassette insertion

The sgRNA expression cassette targeting to intergenic region between fucose and rhamnose operons (Supplementary Table S4) was synthesized and amplified using primers sgRNA-XbaI and sgRNA-XmaI (Supplementary Table S3), then cloned into XbaI-XmaI-digested pMJM100 to generate targeting plasmid pMJM100-rha-sgRNA (Supplementary Table S2). The repair template (Supplementary Table S4) with two 0.5-kb homologous regions was synthesized and cloned into the SalI-ApaI-digested pMJM100-rha-sgRNA generating plasmid prha-sgRNA-RT2 (Supplementary Table S2). The *gusA* gene expression cassette was amplified from *L. gasseri* ADH using primers gusA-SacII-F and gusA-SacII-R and inserted into prha-sgRNA-RT2 at the *SacII* site, generating prha-sgRNA-gusA-RT2 (Supplementary Table S2). All these final plasmid constructs were PCR-screened using PS3 and PS4 primers (pMJM100 based) and sequenced to confirm sequence content.

### Transformation of *Lacticaseibacillus* strains

Electrotransformation of LGG and other *Lcb. rhamnosus* strains was performed following our previously described protocol (Xie et al., 2025), with minor modifications. Briefly, competent cells were prepared as previously outlined and electroporated under identical conditions (2.5 kV, 25 µF, 200 Ω in 2-mm cuvettes). The main adjustments included using 1 μg of plasmid DNA per 100 μL of cells and decreasing the recovery volume to 900 μL of pre-warmed MRS broth. All other steps, including cell preparation, washing, and post-electroporation plating, were carried out as reported previously.

Transformants were screened by PCR and confirmed by Sanger sequencing using gene-specific primers (Supplementary Table S3). For the fucose isomerase gene *fucI*, the primers fucI-check-F and fucI-check-R were used for the chromosomal PCR amplification and sequencing (2.7-kb in the WT and 2.1-kb in deletion mutant). For the CoA synthase gene *acs1*, the primers acs1-check-F and acs1-check-R were used for the chromosomal PCR amplification and sequencing (3.6-kb in the WT and 2.1-kb in deletion mutant). For the insertion of *cam* expression cassette, the primers fucI-check-F and fucI-check-R were used for the chromosomal PCR amplification and sequencing (2.7-kb in the WT and 3.2-kb in insertion mutant). For the insertion of *gusA* expression cassette, the primers fuc-rha-check-F and fuc-rha-check-R were used for the chromosomal PCR amplification and sequencing (2.1-kb in the WT and 4.2-kb in insertion mutant).

### Growth Curve Measurement

The growth of the LGG strain and its mutant *L. rhamnosus* MJM568 and MJM670 was analyzed in sugar-free MRS media (US Biological, Swampscott, MA, USA) containing 2% glucose, fucose, or rhamnose (Sigma, St. Louis, MO, USA). Strains were picked and cultured in MRS broth overnight.

Then the cultures were harvested and washed three times with phosphate-buffer solution (PBS, pH 7.4) and resuspended in same volume of PBS buffer. Cells were inoculated 2% into MRS containing different sugar sources and transferred into a sterile 96-well plate. Each condition was tested in triplicate. The plates were incubated at 37 ℃ without agitation for 24 h in FilterMax F5 Multi-Mode Microplate Reader (Molecular Devices, LLC, Sunnyvale, CA, USA). The absorbance at 595nm (Abs_595nm_) in each well was recorded every 20 or 30 min. The maximum specific growth rate (µ_max_) was defined as the slope of the tangent line at the inflection point of the logistic regression curve and was reported in hours^-1^.

### Assessment of *gusA* Gene Expression Cassette Stability

To evaluate the genetic stability of the *gusA* gene, both plasmid-based (LGG with pMJM100-gusA) and genome-integrated (MJM670) constructs were examined over multiple generations. The *gusA* gene expression cassette was amplified from *L. gasseri* ADH using primers gusA-SpeI-F and gusA-SpeI-R and inserted into pMJM100 at the *SpeI* site, generating pMJM100-gusA (Supplementary Table S2). LGG harboring pMJM100-gusA was cultured 12 hours in MRS broth supplemented with Em (0 generation), while MJM670 was grown 12 hours in MRS broth without antibiotics (0 generation). Then, 0.1% of each overnight culture was then inoculated into 10 mL of fresh MRS broth without antibiotics. This process was repeated every 12 hours for a total of 2 days (∼40 generations). At defined intervals (e.g., 0, 10, 20, 30, and 40 generations), cultures were serially diluted in PBS and plated on MRS agar supplemented with 200 µg/mL X-gluc. The proportion of blue (GUS-expressing) and white (non-expressing) colonies was quantified to assess the retention of *gusA* expression.

### In vivo Mice Study

All animal procedures were approved by the University of Illinois Institutional Animal Care and Use Committee (protocol #23108). Male C57BL/6J mice (8 weeks old, n = 18; Jackson Laboratories, Bar Harbor, ME, USA) were acclimated for 4 days and then randomly assigned to three groups (n = 6 each). Mice were housed individually on wire grids with free access to water and an AIN-93M diet (Research Diets, Inc., New Brunswick, NJ, USA) under a 12-h light/dark cycle. The three experimental groups of mice were administered a single 0.2 mL oral gavage containing either 0.85% NaCl (control), WT LGG (GUS-, 4.6×10^9^ CFU/mL), or MJM670 (GUS+, 3.2 ×10^9^ CFU/mL). Fecal pellets were collected at designated time points and placed into sterile microcentrifuge tubes. For plating analysis, fecal pellets were vortexed vigorously in sterile PBS, followed by serial dilution. Aliquots from each dilution were plated on MRS agar supplemented with X-Gluc and aerobically incubated at 37 °C for colony counting. Plates with colony counts between 25 and 250 were considered within the acceptable range for enumeration and counts outside of this range were excluded from analysis. CFU per gram of feces was calculated based on the dilution factor. Due to the lowest countable dilution being 1:100, the detection limit for this assay was estimated at 2.5 × 10⁴ CFU/g feces.

### Quantitative PCR Analysis

Total DNA was extracted using the PureLink™ Microbiome DNA Purification Kit (Invitrogen, Thermo Fisher Scientific, Waltham, MA, USA) according to the manufacturer’s instructions. DNA concentration was measured with a Nanodrop spectrophotometer (Thermo Scientific, Waltham, MA, USA). LGG was detected using strain-specific primers Lrhamn1 and Lrhamn2 (Brandt & Alatossava, 2003; L. Zhang et al., 2020). qPCR was carried out using our previous reported protocol (Lei et al., 2024) on a Q instrument (Quantabio, Beverly, MA, USA). Reactions (10 μL) contained PowerUp SYBR Green Master Mix (Thermo Fisher Scientific, Waltham, MA, USA), LGG-specific primers, and extracted DNA in the same proportions as described earlier. Cycling conditions included an initial incubation (50 °C, 2 min), denaturation (95 °C, 2 min), and 40 amplification cycles (95 °C for 15 s, 54.5 °C for 15 s, 72 °C for 1 min). Each sample was analyzed in triplicate. Standard curves were generated using serial dilutions of known concentrations of LGG genomic DNA, and results were expressed as colony-forming units per gram of feces (CFU/g). For samples with *Ct* values greater than 35, a fixed *Ct* value of 35—corresponding to the detection limit of 1.1 × 10⁴ CFU/g feces—was assigned to facilitate consistent data analysis.

### Statistical Analysis

Results are presented as mean values ± standard deviation from independent experiments. Differences between groups were evaluated using unpaired *t*-tests and one-way ANOVA, with statistical significance defined as *p < 0.05*. In figures, distinct letters were used to indicate significant different groups. All analyses and graph preparation were performed using GraphPad Prism (GraphPad Software, Boston, MA, USA).

## ACKNOWLEDGEMENTS

Zifan Xie thanks support from the Investment for Growth Program from the University of Illinois, Synthetic Biology for Food and Nutrition Innovation (SynFONI). We would like to express our deep appreciation to Molly Black for her invaluable assistance and unwavering support during the mice experiment, which significantly contributed to the success of our research.

## DATA AVAILABILITY STATEMENT

The authors confirm that the data supporting the findings of this study are available within the article and its supplementary materials.

