## Supplementary information for "Exploiting Endogenous CRISPR-Cas9 System for Functional Engineering of Probiotic *Lacticaseibacillus rhamnosus* GG (LGG)"

Affiliation:

***Corresponding author:**

**Construction of the CRISPR RNA (crRNA)-Based Editing Plasmid**

A double-stranded synthetic crRNA gBlock (Supplementary Table S4) containing a promoter that is the native leader of the CRISPR array of LGG, two partial CRISPR repeats, and a rho-independent terminator was amplified by primers crRNA-gBlock-F/R and cloned into XbaI-HindIII-digested pTRK870 to generate a flexible plasmid, pcrRNA-BsaI, in which self-targeting spacers can readily be cloned (Supplementary Table S2). Conveniently, we designed pBsaI with two BsaI sites between the two partial repeats, allowing flexible and easy insertion of spacers (30-bp) as programmable self-targeting guides, using annealing oligonucleotides with overhang ends compatible with the BsaI-digested plasmid.

To construct *fucI* gene targeting plasmid, the pcrRNA-BsaI plasmid was digested with BsaI and ligated with annealed oligonucleotides carrying overhang ends. The resulting plasmid is a pcrRNA-BsaI derivative containing a spacer to target the gene *fucI*, generating the plasmid pfucI-crRNA (Supplementary Table S2). pfucI-crRNA presents a *SalI-ApaI* restriction site to clone a designed homologous recombination repair template to perform genome editing. For this, two 1-kb homologous regions were fused by overlap extension PCR to obtain the 2-kb repair template to the *fucI* gene. This repair template was cloned into SalI-ApaI-digested pfucI-crRNA and pTRK870 generating the plasmid pfucI-crRNA-RT and pfucI-RT (Supplementary Table S2), respectively. All these resulting plasmids were isolated from *Escherichia coli* MC1061 transformants, checked by colony PCR with PS1 and PS2 primers (Supplementary Table S3) for the presence of the insert, and sequenced to confirm sequence content.

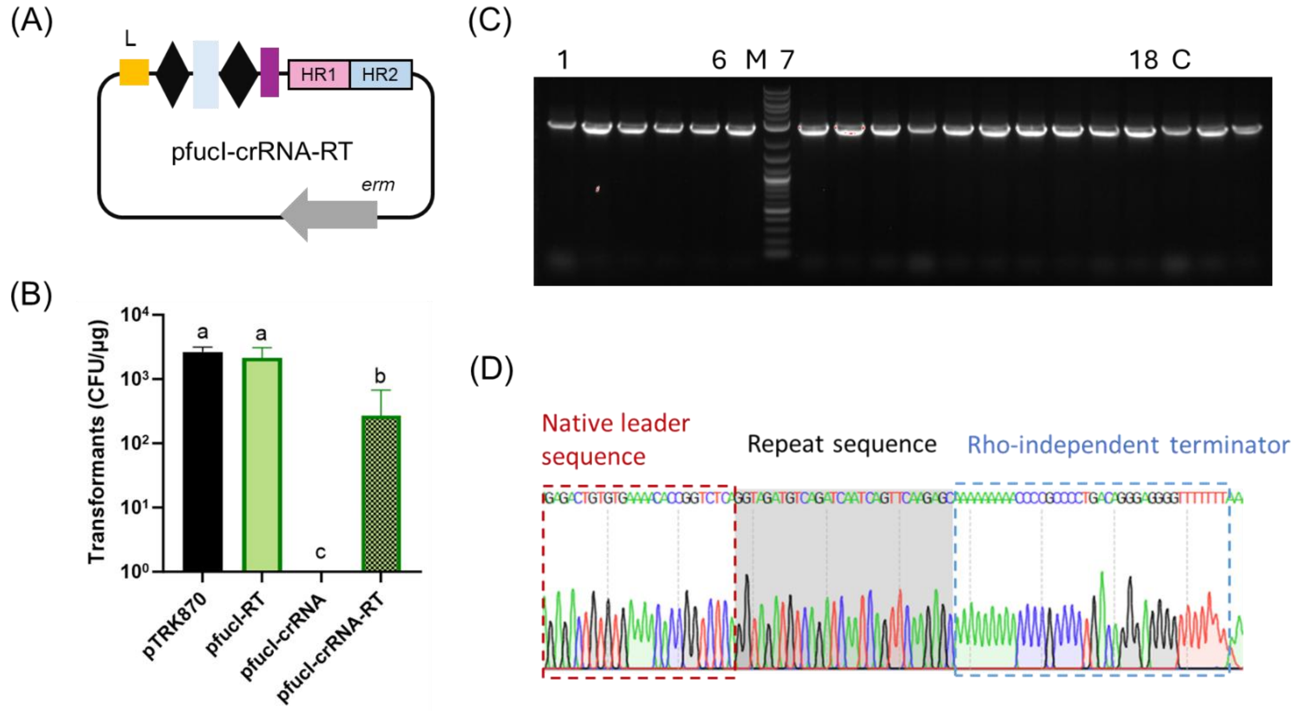


**Supplementary Figure S1. *fucI* gene targeting using artificial crRNA editing plasmid.** (A) An artificial crRNA-based editing plasmid containing the native leader (L, yellow rectangle) of the CRISPR array of LGG as promoter, space targeting to *fucI* gene (blue rectangle), two repeats (native repeat sequence of LGG, black diamonds), a Rho-independent terminator (purple rectangle), and a homologous repair template (HR1 and HR2) were cloned into pTRK870 to generate the artificial crRNA-based editing plasmid, pfucI-crRNA-RT. (B) Transformants of the editing plasmid pfucI-crRNA-RT, the self-targeting plasmid pfucI-crRNA, and the homologous repair template containing plasmid pfucI-RT, with pTRK870 as a control. Distinct letters were employed to denote significant differences ($p < 0.05$). (C) Colony PCR for screening the deletion genotype of 18 randomly picked colonies carrying the editing plasmid, pfucI-crRNA-RT. Expected products sizes corresponding to the WT and the *ΔfucI* strains are 2.7 kb and 2.1 kb, respectively. M: DNA marker; C: WT colony. (D) Sequencing results showed that the space between the repeats was easily looped out from the plasmid during the plasmid replication making a space deficient plasmid.

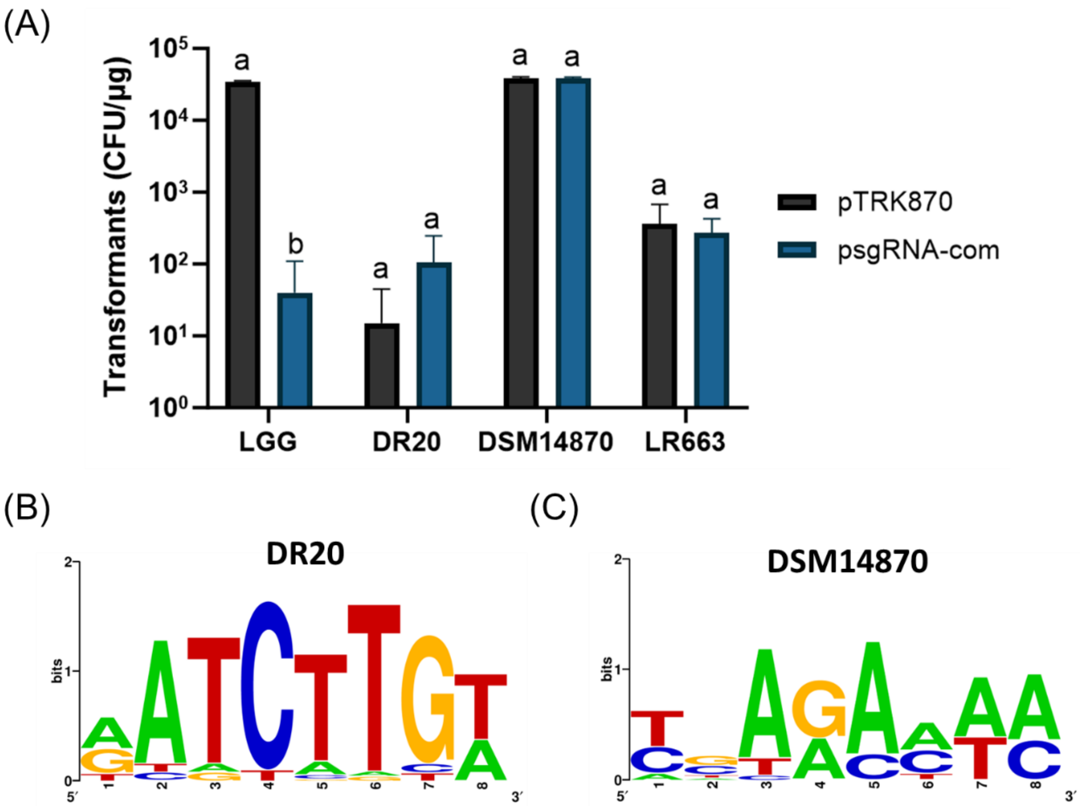


**Supplementary Figure S2. Evaluation of targeting plasmid applicability across different *Lcb. rhamnosus* strains.** (A) Transformation efficiencies of pTRK870 and psgRNA-com in different *Lcb. rhamnosus* strains. Distinct letters were employed to denote significant differences ($p < 0.05$). (B) Predicted PAM sequence for *Lcb. rhamnosus* DR20 based on CRISPR spacer analysis. (C) Predicted PAM sequences for *Lcb. rhamnosus* DSM14870.

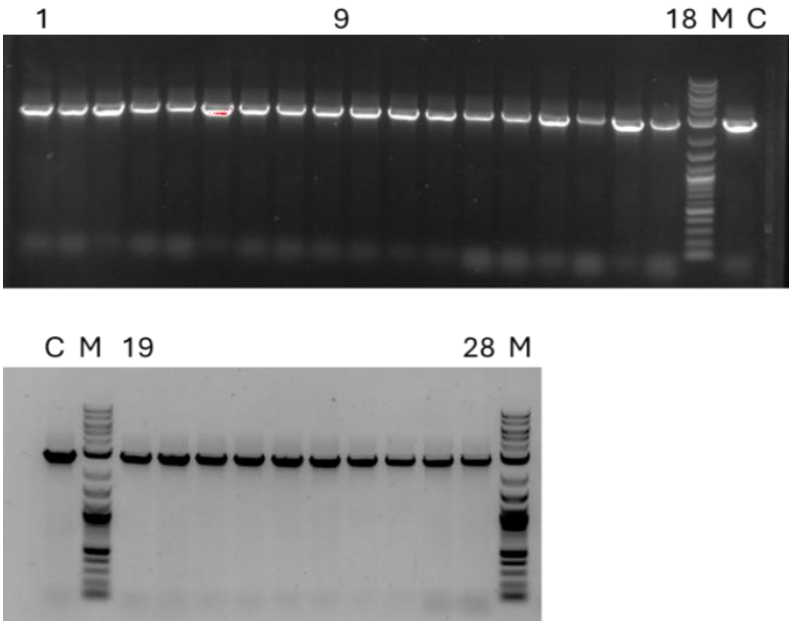


**Supplementary Figure S3. Colony PCR screening of surviving transformants carrying the plasmid pfucI-RT to confirm the deletion genotype.** Expected PCR product sizes for WT and *ΔfucI* are 2.7 kb and 2.1 kb, respectively. C: WT control; M: DNA marker; 1-28: surviving transformants.

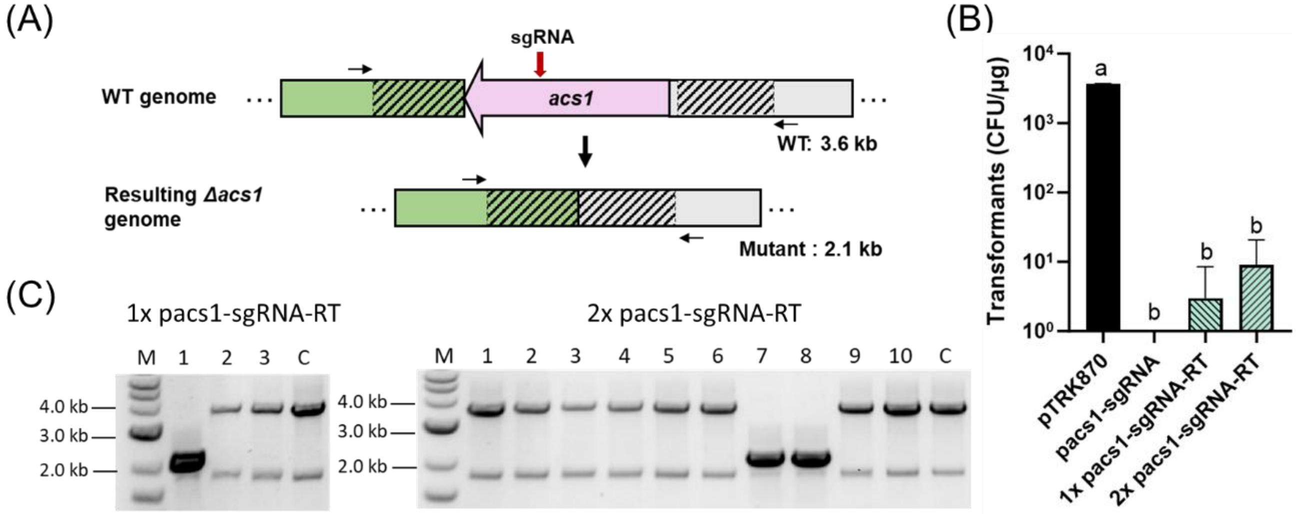


**Supplementary Figure S4. Deletion of coenzyme A (CoA) synthase (*acs1*) by reprogramming the endogenous CRISPR-Cas9 system in LGG.** (A) Schematic representation of the genotypes of the wild type (WT) strain and the *Δacs1* mutant. (B) Transformation efficiencies following electroporation with 1 µg of the control plasmid pTRK870, 1 µg of the self-targeting plasmid pacs1-sgRNA, and either 1 µg (1x) or 2 µg (2x) of the editing plasmid pacs1-sgRNA-RT. Distinct letters were employed to denote significant differences ($p < 0.05$). (C) Colony PCR screening of surviving transformants carrying the editing plasmid pacs1-sgRNA-RT to confirm the deletion genotype. Left: transformation with 1 µg pacs1-sgRNA-RT; Right: transformation with 2 µg pacs1-sgRNA-RT. C: WT control; M: DNA marker; 1-3 and1-10: surviving transformants. Expected PCR product sizes for WT and *Δacs1* are 3.6 kb and 2.1 kb, respectively.

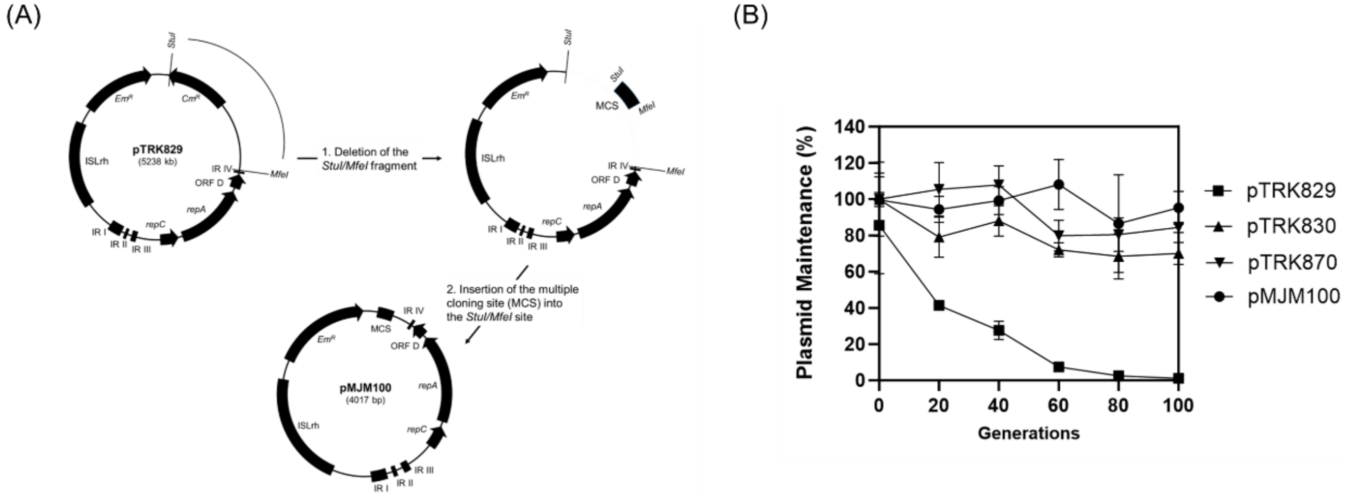


**Supplementary Figure S5. The construction and stability of pMJM100.** (A) Schematic overview of the plasmid pMJM100 construction. (B) Plasmid segregational stability in LGG under non-selective conditions.

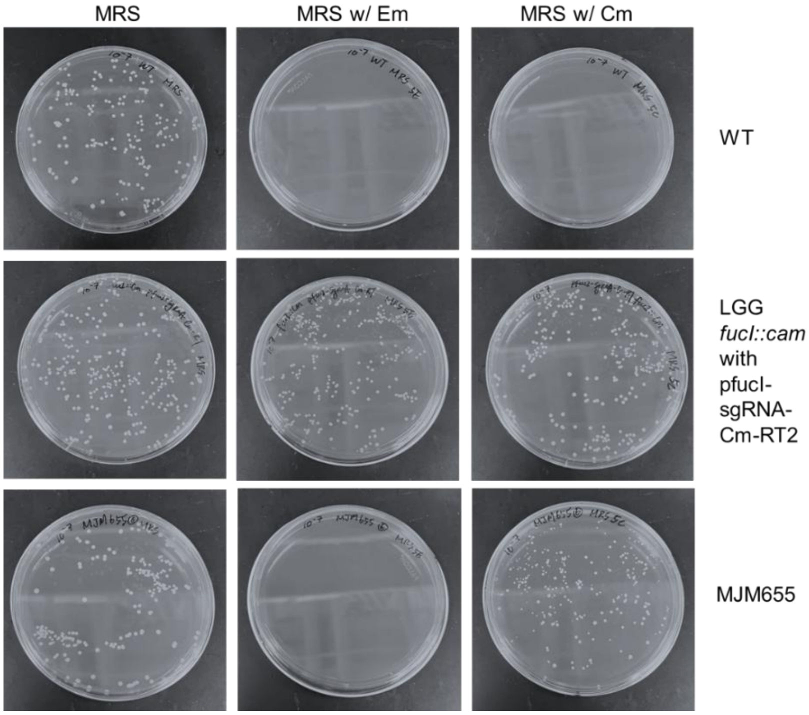


**Supplementary Figure S6. Growth of WT LGG, LGG *fucI::cam* with editing plasmid pfucI-sgRNA-Cm-RT2, and MJM655 strains on MRS plates supplemented with 5 µg/mL chloramphenicol or 5 µg/mL erythromycin.**

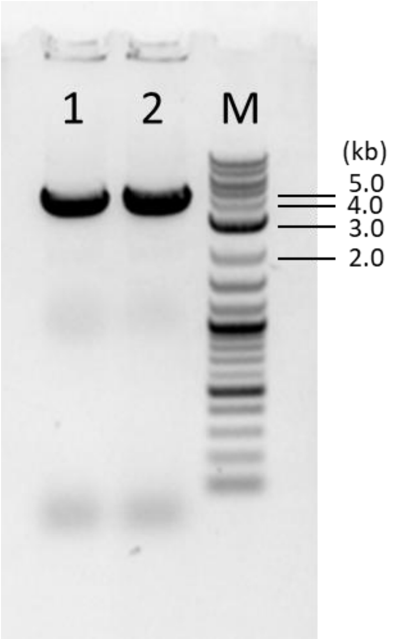


**Supplementary Figure S7. Confirmation of *gusA* insertion mutant purity after re-isolation.** PCR analysis of re-isolated Colony No. 1 showed a single 4.2 kb band corresponding to the correct *gusA* insertion, confirming loss of the wild-type allele observed in the initial screen. M: DNA marker; 1-2: isolated single colonies.

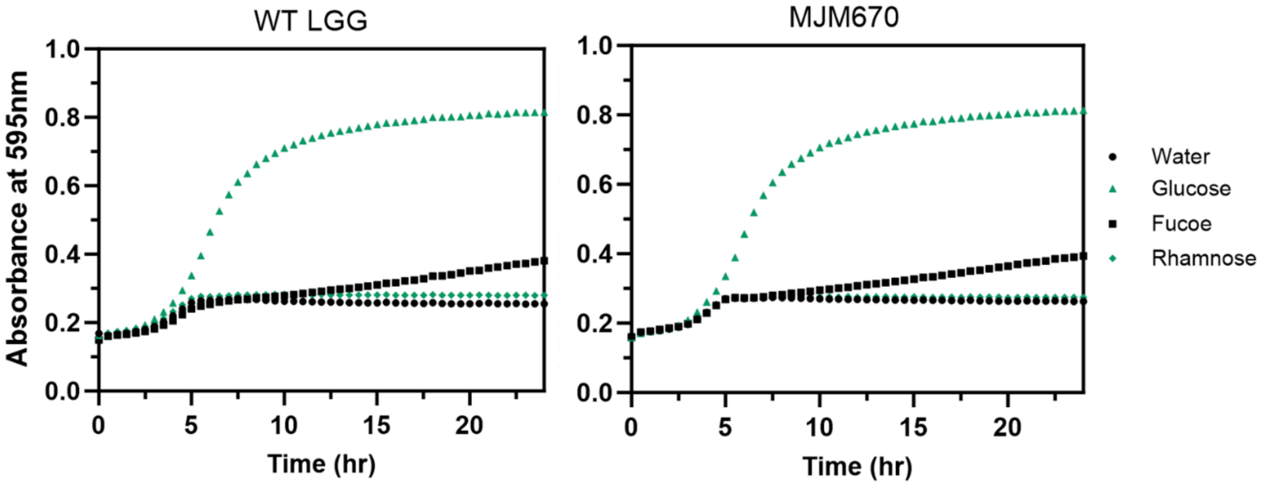


**Supplementary Figure S8. Growth curves of WT and MJM670 strains in MRS supplemented with different carbohydrates.** Growth curves (absorbance at 595nm) of WT (left) and MJM670 (right) strains in MRS with water (circle), 2% glucose (triangle), 2% fucose (square), or 2% rhamnose (diamond).

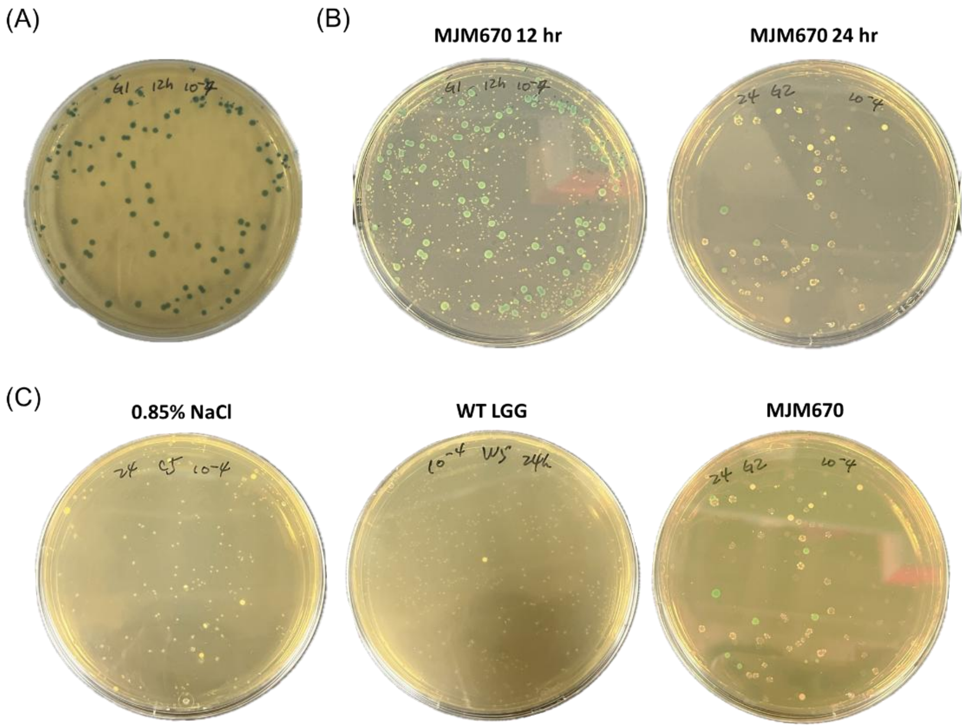


**Supplementary Figure S9. Detection of MJM570 in fecal samples using MRS agar supplemented with X-Gluc.** (A) MJM670 colonies exhibit a distinct blue color against a white background on MRS X-Gluc plates. (B) Fecal samples from mice gavaged with MJM670 were collected at 12 hours (left) and 24 hours (right) after a single gavage and plated on MRS plates with X-Gluc. Blue colonies indicate the presence of MJM670. (C) Fecal samples collected 24 hours after gavage from mice treated with 0.85% NaCl (left), wild-type LGG (middle), or MJM670 (right) were plated on MRS plates with X-Gluc. Only samples from the MJM670 group produced blue colonies, confirming GUS+ activity.

**Supplementary Table S1. Strains used in this study**

| Strains | Description | Sources or references |
|---|---|---|
| *E. coli* MC1061 | Wild-type strain, transformation host | Lab stock |
| *L. gasseri* ADH | Wild-type strain, source of *gusA* gene | Lab stock |
| ***Lcb. rhamnosus*** | | |
| DR20 | Wild-type strain, transformation host | NZDRI |
| LR663 | Wild-type strain, transformation host | Commercial isolate |
| DSM14870 | Wild-type strain, transformation host | Commercial isolate |
| GG, ATCC 53103 | Wild-type strain, transformation host | ATCC |
| MJM568 | *Lcb. rhamnosus* GG mutant with a 640-bp deletion of the L-fucose isomerase *fucI* gene; fucose-negative phenotype. | This study |
| MJM665 | *Lcb. rhamnosus* GG mutant with a 1518-bp deletion of the *acs1* gene. | This study |
| MJM655 | *Lcb. rhamnosus* GG mutant in which a 640-bp deletion of the *fucI* gene was replaced with a 1164-bp chloramphenicol resistance *cam* gene expression cassette; chloramphenicol-resistance ($Cm^R$). The editing plasmid, pfucI-sgRNA-Cm-RT2, is cured in this strain. | This study |
| MJM670 | *Lcb. rhamnosus* GG mutant with a 2077-bp β-glucuronidase (GUS) encoding gene *gusA* (from *L. gasseri* ADH); GUS-positive phenotype. The editing plasmid, prha-sgRNA-gusA-RT2, is cured in this strain. | This study |

**Supplementary Table S2. Plasmids used in this study**

| Plasmids | Description | Sources or references |
|---|---|---|
| pTRK829 | Low copy number *E. coli* and Lactobacilli shuttle vector; $Cm^R$ and $Em^R$. | (1) |
| pTRK830 | Low-copy number *E. coli* and Lactobacilli shuttle vector, derivative of pTRK829; erythromycin-resistance ($Em^R$). | (1) |
| pTRK870 | Cloning vector constructed by inserting a multiple cloning site sequence cloned into pTRK830 at *StuI* site; $Em^R$. | This study |
| **Plasmid interference assay** | | |
| pS1 | Protospacer sequence of spacer 1 from LGG CRISPR array cloned into pTRK870 at *XbaI-HindIII* sites. | This study |
| pPS1AGAAA | Protospacer sequence of spacer 1 and sequence (5'-AGAAA-3') cloned into pTRK870 at *XbaI-HindIII* sites. | This study |
| pPS1TGAAA | Protospacer sequence of spacer 1 and sequence (5'-TGAAA-3') cloned into pTRK870 at *XbaI-HindIII* sites. | This study |
| pPS1GGAAA | Protospacer sequence of spacer 1 and sequence (5'-GGAAA-3') cloned into pTRK870 at *XbaI-HindIII* sites. | This study |
| pPS1AAAAA | Protospacer sequence of spacer 1 and sequence (5'-AAAAA-3') cloned into pTRK870 at *XbaI-HindIII* sites. | This study |
| pPS1ACAAA | Protospacer sequence of spacer 1 and sequence (5'-ACAAA-3') cloned into pTRK870 at *XbaI-HindIII* sites. | This study |
| pPS1TCAAA | Protospacer sequence of spacer 1 and sequence (5'-TCAAA-3') cloned into pTRK870 at *XbaI-HindIII* sites. | This study |
| pPS1AGGAA | Protospacer sequence of spacer 1 and sequence (5'-AGGAA-3') cloned into pTRK870 at *XbaI-HindIII* sites. | This study |
| pPS1AGTAA | Protospacer sequence of spacer 1 and sequence (5'-AGTAA-3') cloned into pTRK870 at *XbaI-HindIII* sites. | This study |
| pPS1AGATA | Protospacer sequence of spacer 1 and sequence (5'-AGATA-3') cloned into pTRK870 at *XbaI-HindIII* sites. | This study |
| pPS1AGAAG | Protospacer sequence of spacer 1 and sequence (5'-AGAAG-3') cloned into pTRK870 at *XbaI-HindIII* sites. | This study |
| **The deletion of *fucI* gene** | | |
| pfucI-RT | Control plasmid containing a repair template with two 1-kb homologous regions for *fucI* gene deletion, cloned into pTRK870 at *SalI-ApaI* sites. | This study |
| pcrRNA-BsaI | Basic plasmid with an artificial crRNA gBlock (leader + two partial repeats + rho-independent terminator) cloned into pTRK870. | This study |
| pfucI-crRNA | Targeting plasmid for the *fucI* gene, constructed by cloning a 30-bp *fucI*-specific spacer (annealed oligonucleotides) into pcrRNA-BsaI. | This study |

| | | |
|---|---|---|
| pfucI-crRNA-RT | Editing plasmid for *fucI* gene deletion, constructed by inserting the 1-kb repair template into pfucI-crRNA at *SalI-ApaI* sites | This study |
| pfucI-sgRNA | Targeting plasmid for the *fucI* gene, constructed by cloning the *fucI* targeting sgRNA expression cassette gBlock into pTRK870 at *XbaI-XmaI* sites. | This study |
| pfucI-sgRNA-RT | Editing plasmid for *fucI* gene deletion, constructed by inserting the 1-kb repair template into pfucI-sgRNA at *SalI-ApaI* sites. | This study |
| **The deletion of *acs1* gene** | | |
| pacs1-sgRNA | Targeting plasmid for the *acs1* gene, constructed by cloning the *acs1* targeting sgRNA expression cassette gBlock into pTRK870 at *XbaI-XmaI* sites. | This study |
| pacs1-sgRNA-RT | Editing plasmid for *acs1* gene deletion, constructed by inserting the 1-kb repair template into pacs1-sgRNA at *PstI-ApaI* sites. | This study |
| **The insertion of *cam* gene** | | |
| pMJM100 | Modified cloning vector derived from pTRK829 by inserting a multiple cloning site sequence at *StuI* and *ApaI* sites, removing the entire *cam* expression cassette and other non-essential regions; $Em^R$. | This study |
| pMJM100-fucI-sgRNA | Targeting plasmid for the *fucI* gene, constructed by cloning the *fucI* targeting sgRNA expression cassette gBlock into pMJM100 at *XbaI-XmaI* sites. | This study |
| pTOPO-fucI-RT2 | Intermediate plasmid for constructing a *cam* gene insertion repair template; a repair template with 0.5-kb homologous regions for *fucI* gene deletion was amplified from pfucI-RT and cloned into a TOPO vector. | This study |
| pTOPO-fucI-Cm-RT2 | Intermediate plasmid for constructing a *cam* gene insertion repair template; the *cam* gene expression cassette from pTRK829 was inserted into pTOPO-fucI-RT2 at the *HindIII* site. | This study |
| pfucI-sgRNA-Cm-RT2 | Editing plasmid for *cam* gene insertion at the *fucI* locus, constructed by cloning the repair template from pTOPO-fucI-Cm-RT2 into pMJM100-fucI-sgRNA at *SalI-ApaI* sites. | This study |
| **The insertion of *gusA* gene** | | |
| pMJM100-rha-sgRNA | Targeting plasmid for the intergenic region between the fucose and rhamnose operons, constructed by cloning the intergenic region targeting sgRNA expression cassette gBlock into pMJM100 at *XbaI-XmaI* sites. | This study |
| prha-sgRNA-RT2 | Editing plasmid for gene insertion at the intergenic region between the fucose and rhamnose operons, constructed by cloning the repair template into pMJM100-rha-sgRNA at *SalI-ApaI* sites. | This study |
| pMJM100-gusA | *gusA* gene expression cassette from *Lactobacillus gasseri* ADH was cloned into pMJM100 at *SpeI* site. | This study |
| prha-sgRNA-gusA-RT2 | Editing plasmid for *gusA* gene (from *L. gasseri* ADH) insertion at the intergenic region between the fucose and rhamnose operons, constructed by cloning the *gusA* gene expression cassette into prha-sgRNA-RT2 at *SacII* site. | This study |

**Supplementary Table S3. Primers used in this study [a]**

| Primers | Sequences (5'-3') |
|---|---|
| PS1 | GCGAAAGGGGGATGTGCTGC |
| PS2 | GCACCCCAGGCTTTACACTTTATGC |
| PS3 | GCAATACAATTGGAGCTCCACCATGG |
| PS4 | AGTCATTAGGCCTTGGTACCGGG |
| sgRNA-XbaI | CGTACCATCGTCTAGATGCGACAAGT |
| sgRNA-XmaI | CGATCCCGGGAAAAAAATGTATGGCC |
| Lrhamn1 | CAATCTGAATGAACAGTTGTC |
| Lrhamn2 | TATCTTGACCAAACTTGACG |
| **Plasmid interference assay** | |
| PS1XbaI | AGTCATCTAGAAAGAAACGATCCCGAGTTTCTGG |
| S1HindIII | AGTCAAAGCTTTAGTAAGCAATCCAGAAACTCGGGATC |
| PS1AGAAAHindIII | AGTCAAAGCTTTAGTTTCTAAGCAATCCAGAAACTCGGGATC |
| PS1TGAAAHindIII | AGTCAAAGCTTTAGTTTTCAAAGCAATCCAGAAACTCGGGATC |
| PS1GGAAAHindIII | AGTCAAAGCTTTAGTTTTCCAAGCAATCCAGAAACTCGGGATC |
| PS1AAAAAHindIII | AGTCAAAGCTTTAGTTTTTTAAGCAATCCAGAAACTCGGGATC |
| PS1ACAAAHindIII | AGTCAAAGCTTTAGTTTTGTAAGCAATCCAGAAACTCGGGATC |
| PS1TCAAAHindIII | AGTCAAAGCTTTAGTTTTGAAAGCAATCCAGAAACTCGGGATC |
| PS1AGGAAHindIII | AGTCAAAGCTTTAGTTTCCTAAGCAATCCAGAAACTCGGGATC |
| PS1AGTAAHindIII | AGTCAAAGCTTTAGTTTACTAAGCAATCCAGAAACTCGGGATC |
| PS1AGATAHindIII | AGTCAAAGCTTTAGTTATCTAAGCAATCCAGAAACTCGGGATC |
| PS1AGAAGHindIII | AGTCAAAGCTTTAGTCTTCTAAGCAATCCAGAAACTCGGGATC |
| **The deletion of *fucI* gene** | |
| fucI-HR1-F | CAAATGCGCAAAGTCACCTGGAAC |
| fucI-HR1-R | CGCATCAAACCCGCCGTCAATTTTCTTCACAGTTCATCAAGCTTATTAG |
| fucI-HR2-F | TGACGGCGGGTTTGATGCGAAATAAGAGTTATC |
| fucI-HR2-R | CCCGGATCTGTTCGCTTGTTG |
| fucI-RT-SalI | TACCGTCGACCAAATGCGCAAAG |
| fucI-RT-ApaI | GTCAGGGCCCCCCGGATCTGTTC |
| fucI-crRNA-F | GGTAGATGTCAGATCAATCAGTTCAAGAGCTGCTTCTTTAACACCGCCAATGGTTGAATCGTCTCA |
| fucI-crRNA-R | TACCTGAGACGATTCAACCATTGGCGGTGTTAAAGAAGCAGCTCTTGAACTGATTGATCTGACATC |
| fucI-check-F | GTCCCGTCAAAATTGGAACAAAGAATGTTG |
| fucI-check-R | CCCAGAGCTTTGTCGAGTCACC |
| **The deletion of *acs1* gene** | |
| acs1-PstI | CATACTGCAGGTTGCGGACCG |

| | |
|---|---|
| acs1-ApaI | GAATGGGCCCTCACTTCACACG |
| acs1-check-F | CTGAGCTGGTGAACGCGTTATGG |
| acs1-check-R | GAACCACACCACCACGGCATTTC |
| **The insertion of *cam* gene** | |
| fucI-RT2-SalI | TACCGTCGACCGTGCGGTTGATGAATGCCATTC |
| fucI-RT2-ApaI | GTCAGGGCCCCGTCTTGCCATATATGACCCCAGG |
| Cm-F | CATCAAGCTTCCACATTTTTAATGTGGTCTTTATTCTTCAACTAAAGC |
| Cm-R | GACTAAGCTTCATCGGCTTTCATAATCTAACAGACAACATCTTCG |
| **The insertion of *gusA* gene** | |
| gusA-SpeI-F | CTGATACTAGTCATTGGCCACTCTCAACATTCCTTTC |
| gusA-SpeI-R | ATGATACTAGTCGCGCCCATAATCAAATAATCGATATC |
| gusA-SacII-F | CTGATCCGCGGTCATTGGCCACTCTCAACATTCCTTTC |
| gusA-SacII-R | ATGATCCGCGGCGCGCCCATAATCAAATAATCGATATC |
| fuc-rha-RT2-SalI | CAGTGTCGACCATAGCACCAACAAGGTGTAACTGTG |
| fuc-rha-RT2-ApaI | CTGAGGGCCCCACTGCCATCCTACCAATCTGGTTG |
| fuc-rha-check-F | CATCCATCAGATCACGAGCAATGATCG |
| fuc-rha-check-R | CGACTGGATGAACAACATTCACCAGTATG |

[a] Introduced restriction sequences are underlined and targeting spacers are double underlined.

**Supplementary Table S4. Sequences of sgRNA expression cassettes and repair templates [a]**

| gBlock | Sequences (5’-3’) |
|---|---|
| crRNA gBlock | GTCATCTAGAGAGACTGTGTGAAAACACCGGTCTCAGGTAGAAAGGTCTCAGGTAGATGTCAGATCAATCAGTTCAAGAGCAAAAAAAAACCCCGCCCCTGACAGGGAGGGGTTTTTTTAAGCTTTCAGT |
| *fucI* targeting sgRNA expression cassette | CGTACCATCGTCTAGATGCGACAAGTAATAAACTAAACAAAACAACTACAAAATATTTCTTTTTGTTTTTCATGATTTTTACACTTCTCTTAGTATGCTTTTGTTATAAGTTAGCACAAAAAAGCAGAAAATAAAAAGTAGAAATAAAAAAAGATGTTTTTTTGCCCATATCTCTATGAAAAAAACTGTGAAATGTGTAAAATATGGATGAAACATTGAATTTAAATGCTTCTTTAACACCGCCAATGGTTGAATCGTCTCAGGTAGATGTCAGATCAATCTCTGACATCTACGAGTTGAGATCAAACAAAGCTTCAGCTGAGTTTCAATTTCTGAGCCCATGTTGGGCCATACATTTTTTTCCCGGGATCG |
| *acs1* targeting sgRNA expression cassette | CGTACCATCGTCTAGATGCGACAAGTAATAAACTAAACAAAACAACTACAAAATATTTCTTTTTGTTTTTCATGATTTTTACACTTCTCTTAGTATGCTTTTGTTATAAGTTAGCACAAAAAAGCAGAAAATAAAAAGTAGAAATAAAAAAAGATGTTTTTTTGCCCATATCTCTATGAAAAAAACTGTGAAATGTGTAAAATATGGATGAAACATTGAATTTAAATCTTGATCCAGATATCCGAGGTCACCAGTTGTCTCAGGTAGATGTCAGATCAATCTCTGACATCTACGAGTTGAGATCAAACAAAGCTTCAGCTGAGTTTCAATTTCTGAGCCCATGTTGGGCCATACATTTTTTTCCCGGGATCG |
| Repair template for *acs1* gene deletion | CATACTGCAGGTTGCGGACCGAATCATCTAGCTGGTTTTAATTGGAATCGCTGACCAGGGTTGTCTTCATATCAACCTTCGTCGTGATAACCCCATCCTGATAAAACACATCGGCAATGGCGTTTTCTTCTGCCACAATGCTACTGTTGGCACTAATCAGCGTCATGGAATAGGTGCGCCGTTCTACCATCTTTTTAATCACTTTATCGGAGAGATTGAGGGTTTTACTGAGCATCGCAATGAGTTGCGTATGATGGCTATTGGCCCAAGTCATATCGTCATTTAGTTCAGTCGTCAGTAACTTACTTAATGATGTATGGGTCTTGGCATAGCTCTGGGTTGAAATCAGGAAATCACGGTTCTTGGTCAAACCGGTGCCGTTGGTCAACAGCTTAGCACCTTGATTGACTTGGGCCGTTGACGTATATGGATCCCAGGTAACCCATGCATCGACACTACCTTTAGCAAATGCAACCGAGGCCGAGCTTTGATCCATGTTCACCAGTTTCACATCACTGGTGCTGAGTCCGGCTTTCTTGAGCGCCTGGATGATAAGGTACTGGGCGGCAGTGCCTTTTTGATACGCAATGGTCTTGCCCTTTAATTGCTCAAGACTGGTGATGCCGCTATTTTTACCAACCAAAATACCACTACCATATTCTTTGGTGGCCCCAGCGGCAATTAACGCAATATCCGTGCCAGTTGCTTTGGCAGTCACAGGCGGTGTATCCCCGACCCGCGCATAGTCGATGGCACCGCTTTTAAGTGCCGTCATCAAGGCTGCTCCGTCAGAGAACTCCTTGAACTCAACCTGATAGCCTTTTGCTTTCAACTTTTTGGCTAATTCGCCACGCTGCCGGGCAATGTCGACAGGATCGGCTTTTTGATAGCCAATCGTGACGGTCTCAAGGTTGCTCGATGCCGTGGCGGTTTGCGTATAACCGTAAACCGCCACGCTCAGCCAAGCGACTAAGCCCACTAACAATAAAACCAGTCGTAACTTACGCTTCGCTCATGATTCAGGACTCCTTTCGTAGCTAACAACAATGACTGCGGACGGCCAATGAGCCATCCAAACGGGGAAACTTTGTAACAAAACCAAGCGATGGCAAAGGAAATGCCGAGAAT |

| | |
|---|---|
| | GCTGAGATAACCCAGCGGCAACAGTAGCATTAACATGAGATTACTAAGATGAACCAGTTCCA<br>AGACACCGCCTAATAGCGTGATGGCAATGGTTTGGCAAAGATAGACACCAAACGAAACTTGT<br>GCGCCATGACCAACCAAACGAGCAAACCAGTGCGTTGGCTGTTTGGCGTGCCACCTGGCAT<br>ACTTTTGACCAAGGATGGCGACAAAGGCGATCATCACCACGTCATATACCAACATATACGGCT<br>GATGAGGTGAAACCGCAGCACTGTGAGTCAGGCCAAGAATCTGTTTGTTGAAGCCATAAAG<br>ACCAACTGTTCCTAAGCTGAGAAACAAGGTGCTCCAGCCGATCAGGCGCTGATGGGACATTA<br>ACCAGGTTTTAATCGCTTGGGCGTGTAGGCTGAATAAACCACCGGCAATGAAGAAGAATTGG<br>TAGGTCACAACATTGACGCCATATGCTCTAAACCACCATAACCAGTGGCTCGTGTCGATATGC<br>GGCAGCCCATACTTGATGCCTGCGACAATGAGTAACTGTAAAGTAAAACTAGCTAGCAACAA<br>GGCGCCGTGATGCTTTTCCGTTTTTTTGAATAGCCAAAGCATCGCCGGAAATAACAGATACAG<br>TTGTAAGGTTACCAAGACGTAATACAGATAAAATTGATCACCATGCAATAAGGCACTGAGATA<br>ATCCGGCCAGAAGTTGGCAACGGTGAAGCCGCCAATGCCCAGTAGCACGGCACTGATCATTA<br>AGATCGTGTTCCACAAGACATAGGGAACGCCAACTGTCGTGAAACGTTTTTGAAGAACCG<br>CGGCCAGTTACGTTGCCGGGAGTAGTAGACCATCGTCAGCACCAGCCCTGTCATGAACATGA<br>AGCCCATGCGTGTGAAGTGAGGGCCCATTC |
| The intergenic region targeting sgRNA expression cassette | CGTACCATCGTCTAGATGCGACAAGTAATAAACTAAACAAAACAACTACAAAATATTTCTTTT<br>TGTTTTTCATGATTTTTACACTTCTCTTAGTATGCTTTTGTTATAAGTTAGCACAAAAAAGCAG<br>AAAATAAAAAGTAGAAATAAAAAAAGATGTTTTTTTGCCCATATCTCTATGAAAAAAACTGTG<br>AAATGTGTAAAATATGGATGAAACATTGAATTTAAAGAAGGCAATTGGTGGGCTTTTTGGGTT<br>GGAGTCTCAGGTAGATGTCAGATCAATCTCTGACATCTACGAGTTGAGATCAAACAAAGCTT<br>CAGCTGAGTTTCAATTTCTGAGCCCATGTTGGGCCATACATTTTTTTCCCGGGATCG |
| Repair template for *gusA* insertion | CACTGCCATCCTACCAATCTGGTTGCCATGTCCTTTACGCTGCCGCTGGAAGAGAAGCTCTTT<br>TCCCGCATTTTGTGGAAGATGCAGGCTGAATCCATTGTGGTCTTCCCCGAAGGCATCGGCGTC<br>TTGCCATATATGACCCCAGGCACCAACGAAATCGGTCAGGCAACCGCCCAAAAAATGGCCGA<br>TTTCCGAATCGTCATGTGGCCGCAACACGGAATCTTCGGCGCCGGCGATTCCATTGATGAAA<br>CGTATGGCTTAATTGAAACCGTTGAAAAAGCGGCGACAATTTACACCGCTATTCAGGCTCAA<br>GGCGGCCGCATCATCAACGAAATCACGGATGAAAATCTGGAACAACTGGCCAAACGGTTTG<br>ACTTAACGCCTAACCCAGCCTTTCTTCACGGTGATTCACTCATCAGTCAGGTAACGCTAAAA<br>GTGTAACGTTATCTGCTCATGGTAGGATACCTCCTAAACACACTCTAAAGAGCCCACTGATTG<br>TTCCATGGCCGCGGGCTATACATGGTTTCGCCGATGAGGTGGTATAGGAACTTTTATGTGTCT<br>GGGCTAATAAGCTTGATGAACTGTGAAGAAAATTGACGACCTTTTTTGAAGAAAAGCCGTTA<br>ATCGTGCAATCTATTGGCGCGTTGAATGAAAGCGCTTATTATTATAGCCGTAGTCAGCTGATGA<br>ACAAAATATTGTTAGATTTGGTAGATATCAAACTTGGAGGCAGAATAAATGACAAACGTGAC<br>ATTTCCTAAAATTGGTATTCGTCCGACGATTGATGGACGCCGGCGTGGTATTCGTGAATCACT<br>AGAAGAACAGACTATGAACATGGCCCATTCAGTTGCCAAGCTTTTAGAATCAACCCTTAAAT<br>ATCCAGATGGTTCACCTGTTCAGACTGTTATTTCTGATTCAACCATTGGCGGTGTTAAAGAAG |

| | |
|---|---|
| | CAGCAATGTCTAAGGAAAAATTTGATCGCGCCGGAATTTGTGGAACCATCACAGTTACACCT<br>TGTTGGTGCTATG |

[a] Introduced restriction sequences are underlined and targeting spacers are double underlined.
